# Calcium signaling recruits Rab27a and the exocyst complex to promote small extracellular vesicle release in skeletal muscle cells

**DOI:** 10.64898/2026.10.22.761013

**Authors:** T. Nelson Luke, Jeremy Boulestreau, Noemi Polgar

## Abstract

Extracellular vesicles (EVs) are key mediators of intercellular communication with emerging therapeutic potential. Small EVs (∼50-150 nm) released from skeletal muscle (SkM) following exercise improve insulin sensitivity and glucose tolerance in animal models, yet the molecular mechanisms governing their secretion remain poorly understood. Ca^2+^ signaling has been shown to regulate numerous membrane trafficking events ranging from vesicle transport, membrane repair, and exocytosis of cellular cargo. We investigated how Ca^2+^ signaling influences the intracellular trafficking and membrane delivery mechanisms necessary for EV release using an *in vitro* SkM cell model. We found that elevated intracellular Ca^2+^ stimulates EV secretion through a mechanism requiring activity of the octameric exocyst complex. Using proximity ligation assays, co-immunoprecipitation, structural modeling and proximity-dependent proteomics, we identified dynamic coordination between the exocyst complex and the small GTPase, Rab27a, during Ca^2+^-stimulated EV release. We found this coordination is modulated by Ca^2+^ signaling and perturbation of exocyst function. We then used direct binding assays to provide a structural basis for the proximity observed between GTP and GDP-loaded Rab27a and exocyst subunits. Additionally, proximity-dependent proteomics revealed that Ca^2+^ stimulation promoted the association of specific exocyst subunits with GTP-competent Rab27a, suggesting a nucleotide-state-dependent organization of this trafficking interface in living cells. Finally, we adapted a real-time reporter of EV exocytosis in SkM cells and identified Ca^2+^-responsive kinases that modulate EV secretion. Collectively, our findings support a model in which Ca^2+^ signaling reorganizes interactions between Rab27a and the exocyst to promote intracellular trafficking necessary for EV secretion in skeletal muscle cells.

## INTRODUCTION

Skeletal muscle (SkM)-derived extracellular vesicles (EVs) have been shown to exert significant metabolic effects by facilitating organ-organ communication between SkM and other tissues (1, 2). In addition to promoting glucose tolerance, angiogenesis, and extracellular matrix remodeling in cardiac tissue (3–5), SkM-derived EVs have also been identified as potential clinical biomarkers of muscle health that can indicate both healthy and pathological states (6, 7). These diverse functions highlight the broad therapeutic and diagnostic potential of SkM EVs for regenerative medicine and treatment of metabolic diseases and muscle-wasting disorders. Despite the potential utility of these SkM-derived EVs, there is a fundamental gap in our molecular understanding of how SkM regulates EV biogenesis and secretion.

Intracellular Ca^2+^ signaling regulates multiple membrane trafficking events, including secretory vesicle transport, cytoskeletal remodeling, and membrane fusion. One of the important small EV classes implicated in offering therapeutic benefits includes exosomes. Exosome release requires trafficking of multivesicular bodies (MVBs) that enclose the exosome precursors to the plasma membrane followed by membrane fusion. Ca^2+^-responsive trafficking machinery is likely to play a critical role in regulating the trafficking and fusion machinery involved in this process. We set out to characterize the molecular coordination driving small (∼50-150nm) EV release following calcium Ca^2+^ stimulation in skeletal muscle cells.

EV biogenesis and secretion require coordinated activity among endosomal sorting machinery, cytoskeletal transport systems, membrane tethering, and fusion complexes (8, 9). Rab GTPases direct MVBs to engage with secretory networks. The destination of the Rab GTPase-vesicle complex is largely determined through interactions with specialized proteins that bind these small GTPases, known as effectors. Rab identity, GTP-bound state, and post-translational modifications facilitate the relationship between Rab GTPases and their effectors and promote selective transfer of vesicles to their destinations within subcellular compartments (10, 11).

The exocyst complex is an evolutionarily conserved octameric tethering complex that facilitates the targeted docking of intracellular vesicles to the plasma membrane. The exocyst has been shown to interact with multiple Rab and Ral GTPases, and is thought to act as a scaffold that coordinates vesicle docking with Soluble N-ethylmaleimide-Sensitive Factor Attachment Proteins receptors (SNARE)-mediated membrane fusion (12–17). Regulation of exocyst activity requires dynamic assembly and disassembly of the 8-subunit complex, which is further organized into 2 separate subcomplexes (18). Various stimuli, including phosphorylation of the exocyst subunits, can influence the assembly of the exocyst and thereby affect membrane delivery and tethering of membrane-bound vesicles (19–21). Regulators of vesicular trafficking, such as Rab and Ral GTPases, have been shown to recruit exocyst subunits and drive fusion between vesicular and plasma membranes (22–24). Although early studies of the exocyst revealed that all exocyst subunit genes (EXOC1-EXOC8) are expressed in SkM (25, 26), almost no prior studies have investigated the biological functions of the exocyst in SkM. Our recent work demonstrated that the exocyst facilitates targeted docking of intracellular vesicles carrying glucose and fatty acid transporters to the plasma membrane in response to insulin and Ca^2+^ signaling in SkM cells (27). In addition to facilitating the exocytosis of fuel transporters, both genetic knockout and chemical inhibition-mediated modulation of exocyst function significantly alter the secretion dynamics of various vesicular cargo, including EVs (28, 29). The regulated membrane delivery of fuel transporter-containing intracellular vesicles and exosome-precursor carrying MVBs share many components, including small GTPases, cytoskeletal fibers, molecular motors, as well as the SNARE complex responsible for membrane fusion. Based on these observations, we hypothesized that Ca^2+^ signaling in skeletal muscle cells promotes MVB membrane trafficking and subsequent EV release through dynamic protein-protein associations between Rab GTPases and the exocyst complex.

## RESULTS

### Validation of an *in vitro* model for intracellular calcium flux

We aimed to mimic the rapid and robust changes in Ca^2+^ levels observed in SkM cells upon contraction signaling by establishing a cell culture model using murine C2C12 myotubes and the Ca^2+^ ionophore, ionomycin. We treated these differentiated myotubes with a range of ionomycin concentrations and probed for changes in intracellular Ca^2+^ using the ratiometric Ca^2+^ indicator dye, FURA 2-AM. We found that treatment with 1 μM ionomycin introduced intracellular Ca^2+^ increase in a nearly linear pattern over 30 minutes as opposed to higher concentrations of ionomycin that introduced a sharp rise in intracellular Ca^2+^ levels within 5 minutes of incubation (**Fig. 1A**). Treatment with 10 μM ionomycin produced a larger increase of intracellular Ca^2+^ than the lower concentrations tested and was also comparable to the kinetics and peak achieved with 20 μM ionomycin treatment. Based on these measurements, 10 μM ionomycin was selected to stimulate Ca^2+^-signaling in differentiated C2C12 myotubes for our subsequent experiments. Next, we assessed the viability of ionomycin-treated myotubes utilizing ethidium homodimer-1 (EthD-1), a nucleic acid stain that can selectively label dead and dying cells through activated fluorescence when bound to exposed chromatin. Differentiated C2C12 myotubes incubated with 10 μM ionomycin did not show any significant changes in viability compared to those treated with 0 μM ionomycin (10 mM EtOH as vehicle) after 15 minutes based on EthD-1 fluorescence (**Fig. 1B**). Both conditions showed significantly lower levels of EthD-1fluorescence than myotubes treated with 5 M EtOH (30% EtOH) as a positive control for cell death. We present that our treatment conditions (10 µM ionomycin for 15 minutes) resulted in a high influx of intracellular Ca^2+^ as well as preserved myotube viability.

**Figure 1.**
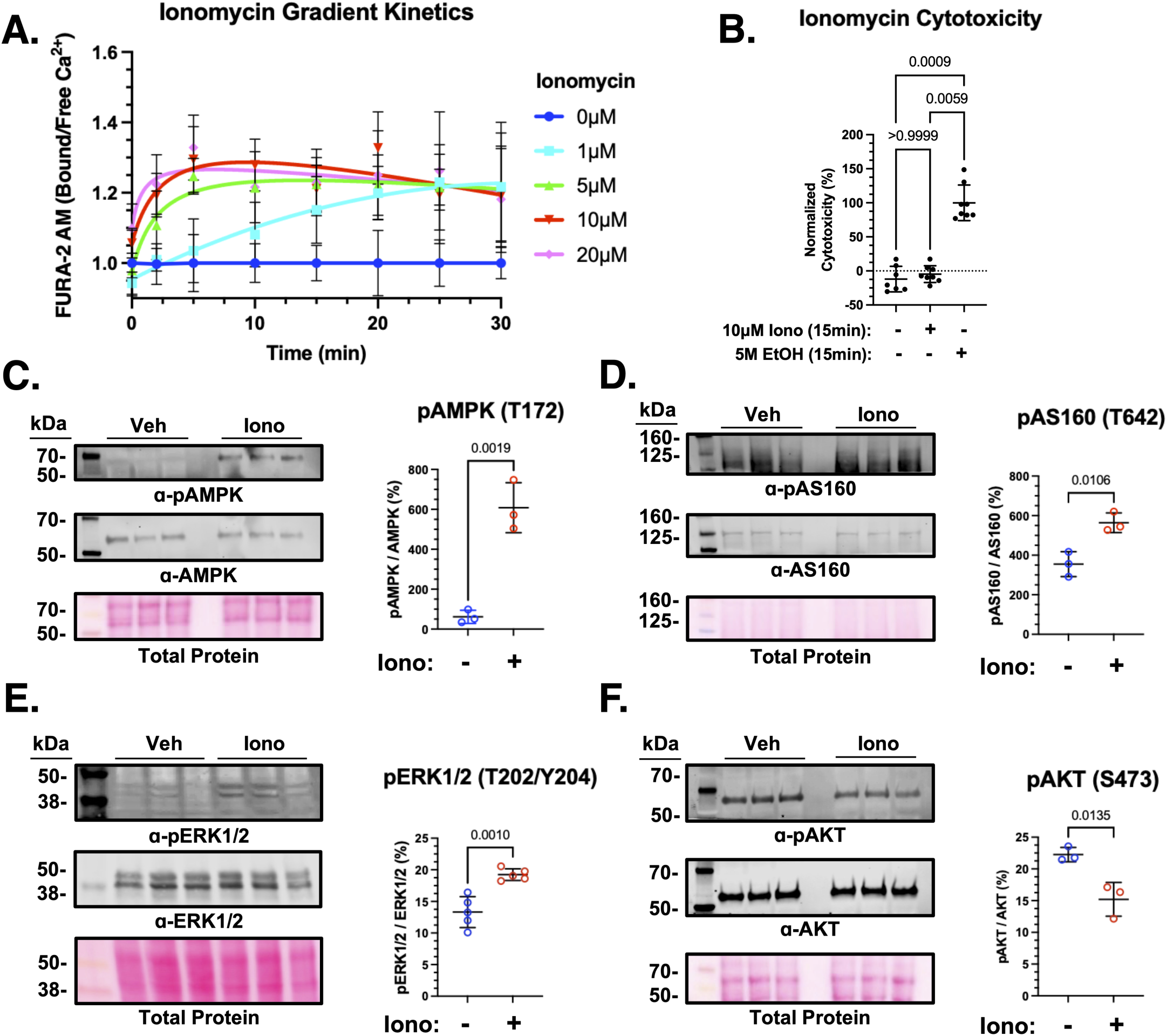
Treatment of differentiated C2C12 myotubes with the calcium ionophore ionomycin as a model of intracellular calcium flux. A. Visual representation of the ratio of intracellular (bound) vs. extracellular (free) Ca^2+^ levels via measurements of the ratiometric dye, FURA2-AM during treatment of C2C12 myotubes with ionomycin. B. Comparison of normalized cytotoxicity (%) in C2C12 myotubes treated with vehicle (EtOH, 10 mM, 15 min), ionomycin (Iono, 10 μM, 15 min), or 5 M EtOH (15 min). One-way ANOVA revealed a significant difference among treatment groups (F(2,64) = 331.3, p < 0.0001), followed by Tukey’s multiple-comparisons test. Cytotoxicity did not differ between vehicle and ionomycin treatment (p = 0.9701), whereas 5 M EtOH treatment resulted in significantly greater cytotoxicity compared with vehicle and ionomycin treatment (p < 0.0001 for both comparisons). Data are shown as mean ± SD; n = 21–23 wells per treatment pooled across three independent experiments. Western analysis of C2C12 lysates following treatment with vehicle (EtOH) or ionomycin. C. Quantification of phosphorylated AMPK (Thr172) relative to total AMPK following vehicle or ionomycin treatment (p = 0.0019). D.Quantification of phosphorylated AS160 (Thr642) relative to total AS160 (p = 0.0106). E. Quantification of phosphorylated ERK1/2 (Thr202/Tyr204) relative to total ERK1/2 (p = 0.0010). F. Quantification of phosphorylated AKT (Ser473) relative to total AKT (p = 0.0135). Unpaired t test. Data are representative of three independent experiments (n = 3) and are shown as mean ± SD.

To determine if ionomycin treatment induced activation of major Ca^2+^-signaling pathways related to actin reorganization, membrane remodeling, and recruitment of vesicular trafficking machinery, we compared phosphorylation levels of the AMP-activated protein kinase (AMPK), extracellular signal-regulated kinase 1/2 (ERK1/2), the GTPase activating protein (GAP), AKT substrate 160 (AS160) and Protein Kinase B (AKT) in vehicle and ionomycin-treated C2C12 myotubes (**Fig. 1C-F**). Myotubes treated with the Ca^2+^ ionophore displayed significantly higher levels of AMPK phosphorylation on Thr172 when compared to vehicle control. We also detected significantly higher levels of phosphorylated AS160 compared to total AS160 levels in response to ionomycin treatment versus vehicle-only control. Ionomycin treatment of differentiated C2C12 myotubes also led to an increased phosphorylation of ERK1/2 at Thr202/Tyr204, while phosphorylation of Protein Kinase B (AKT) at Ser473 significantly decreased in response to ionomycin treatment. Through treatment of C2C12 myotubes with ionomycin, we demonstrate a fast and sustained rise in intracellular Ca^2+^ levels, preserved cell viability and dynamic regulation of kinases associated with Ca^2+^-sensing and handling.

### Exocyst inhibition impairs the effect of ionomycin treatment on skeletal muscle cell EV release

Upon observing that ionomycin treatment induced a rise of both the intracellular Ca^2+^ levels and the activity of Ca^2+^-responsive kinases in our *in vitro* model system, we interrogated the effect of ionomycin treatment on EV release in these cultured skeletal muscle cells. First, we treated differentiated C2C12 myotubes with vehicle or ionomycin and isolated small (∼50-150 nm) EVs. As differentiated myotubes present a multi-nucleated morphology, which can make conventional calculations of EV per cell challenging, we verified that the number of cells producing EVs was consistent between treatments using DAPI quantification of the number of nuclei/image (**Supp. S2)**. Per MISEV standards (30), we visually validated EV morphology and approximate size of the enriched EVs via transmission electron microscopy (TEM) (**Fig. 2A, Supp. S1**) and confirmed the presence of EV markers (CD9, CD81, TSG101) in the EV samples compared to the C2C12 whole cell lysate (WCL) (positive for ER marker, Calnexin) with Western blot analysis (**Fig. 2B**). Using Tunable Resistive Pulse Sensing (TRPS, Izon Sciences), we observed a significant rise in small (∼50-150 nm) EVs derived from conditioned cell medium taken from ionomycin-treated C2C12 myotubes ((3.76 ± 1.22) × 10^10^ EV/mL) vs vehicle-treated C2C12 myotubes ((4.73 ± 2.01) × 10^9^ EV/mL) (**Fig. 2C**). These results confirmed that we can stimulate small EV release in our differentiated C2C12 myotubes with ionomycin treatment.

**Figure 2.**
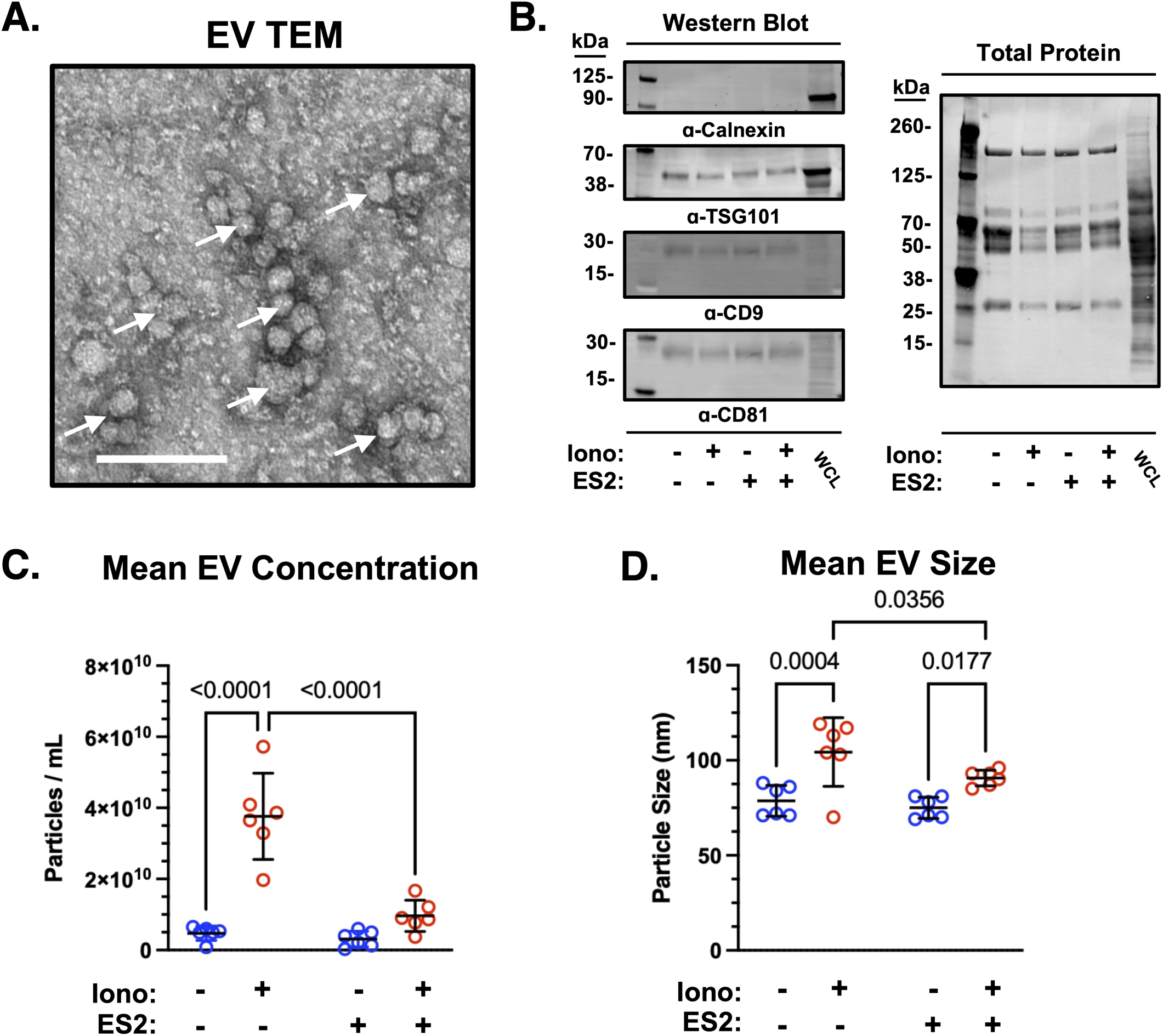
Characterization of small EVs collected from differentiated C2C12 myotubes following ionomycin treatment. A. Representative transmission electron microscopy (TEM) image of isolated EVs. Arrows indicate EVs; scale bar, 200 nm. B. Western blot analysis of isolated EVs and whole-cell lysate (WCL) from C2C12 myotubes probed for the EV markers CD81, CD9, and TSG101 and the endoplasmic reticulum marker calnexin. C–D. EV concentration (C) and particle size (D) following treatment with vehicle (EtOH) or ionomycin in the presence of DMSO or Endosidin 2 (ES2), measured by tunable resistive pulse sensing (TRPS) over an approximately 50–250 nm range. For EV concentration, two-way ANOVA revealed significant effects of ionomycin treatment (F(1,20) = 52.99, p < 0.0001), ES2 treatment (F(1,20) = 29.91, p < 0.0001), and the ionomycin × ES2 interaction (F(1,20) = 23.69, p < 0.0001), followed by uncorrected Fisher’s LSD multiple-comparisons tests. Ionomycin significantly increased EV concentration in DMSO-treated samples (p < 0.0001) but not in ES2-treated samples (p = 0.1035). ES2 significantly reduced EV concentration following ionomycin treatment (p < 0.0001) but not vehicle treatment (p = 0.6748). For EV size, two-way ANOVA revealed a significant effect of ionomycin treatment (F(1,20) = 23.23, p = 0.0001), but no significant effect of ES2 treatment (F(1,20) = 4.086, p = 0.0568) or ionomycin × ES2 interaction (F(1,20) = 1.360, p = 0.2573), followed by uncorrected Fisher’s LSD multiple-comparisons tests. Ionomycin significantly increased EV size in both DMSO-(p = 0.0004) and ES2-treated samples (p = 0.0177). ES2 did not significantly alter EV size under vehicle conditions (p = 0.5522) but significantly reduced EV size under ionomycin conditions (p = 0.0356). Each data point represents an independent experiment (n = 6). Data are shown as mean ± SD.

Docking of MVBs to the plasma membrane is a necessary step for the release of the exosome class of small EVs. We hypothesized that, given that the exocyst complex is known to be responsible for membrane tethering of secretory vesicles prior to cargo exocytosis, inhibition of the exocyst complex would challenge the release of these small EVs in C2C12 myotubes. To perform pharmacological inhibition of exocyst function, we utilized Endosidin 2 (ES2), a small molecule that binds the EXOC7 exocyst subunit and interrupts holocomplex assembly dynamics (31). After pre-treating the C2C12 myotubes with ES2 (30 μM, 30 min) (32), we exposed them to vehicle or ionomycin in the presence of ES2 and collected small EVs from the conditioned cell medium. We found that samples isolated from ES2-treated C2C12 myotubes were not significantly different in their mean concentration ((3.10 ± 2.22) × 10^9^ EV/mL) from untreated controls. However, treatment with ionomycin in the presence of ES2 failed to increase EV concentrations as ionomycin alone did ((9.63 ± 4.39) × 10^9^ EV/mL) (**Fig. 2C**). Of note, when we measured the size of vesicles released under these treatment conditions, EVs collected from ionomycin-treated myotubes had a significantly increased average size compared with EVs collected from vehicle-treated myotubes under DMSO conditions (104.3 ± 18.1 nm versus 78.7 ± 8.1 nm, respectively). This increase in EV size was also observed following ionomycin treatment in the presence of ES2 (90.7 ± 4.1 nm versus 75.0 ± 5.6 nm for ES2-treated vehicle controls). Notably, ES2 significantly reduced the average size of EVs released following ionomycin treatment compared with ionomycin treatment alone (90.7 ± 4.1 nm versus 104.3 ± 18.1 nm), whereas ES2 did not significantly alter EV size under vehicle conditions (**Fig. 2D**). Together, these findings establish that ionomycin treatment triggers small EV release in our in vitro model system and that exocyst inhibition significantly reduces the increase in EV particle concentration associated with ionomycin stimulation.

### Ca^2+^ signaling is linked with increased exocyst subunit proximity

Although the exocyst complex is composed of 8 separate subunits, the rate-limiting association of all 8 subunits depends on an upstream polymerization of two separate subcomplexes. These subcomplexes, each composed of 4 distinct subunits, assemble to generate the octameric exocyst holocomplex (**Fig. 3A**). Exocyst subcomplex assembly has been characterized during various stages of vesicle trafficking, ranging from autophagosome formation to post-Golgi secretory vesicle tethering, yet the state of exocyst subcomplex assembly during increased Ca^2+^ signaling in skeletal muscle cells remains elusive. We used proximity ligation amplification assays (PLA) to detect and quantify associations (≤ 40 nm) between subcomplex II subunits EXOC5 and EXOC7 during treatment with vehicle or ionomycin in differentiated C2C12 murine myotubes. We quantified fluorescent PLA puncta/nuclei that indicate the proximity of EXOC5 and EXOC7 and found that treatment with ionomycin was associated with significantly higher ratios of PLA puncta/nuclei, while the vehicle-treated cells displayed significantly less PLA signal (**Fig. 3B**). We also implemented two negative control groups with either an EXOC5 and Histone H3 (H3.3A) antibody pair or only a single antibody against EXOC5 (**Fig. 3C**) to define the baseline level of off-target/stochastic events in the assay. The increase in PLA puncta/nuclei observed in ionomycin-treated myotubes versus vehicle control suggests that ionomycin treatment in C2C12 differentiated myotubes triggers an increase in proximity between EXOC5 and EXOC7 subunits, an upstream event required for exocyst assembly and tethering activity.

**Figure 3.**
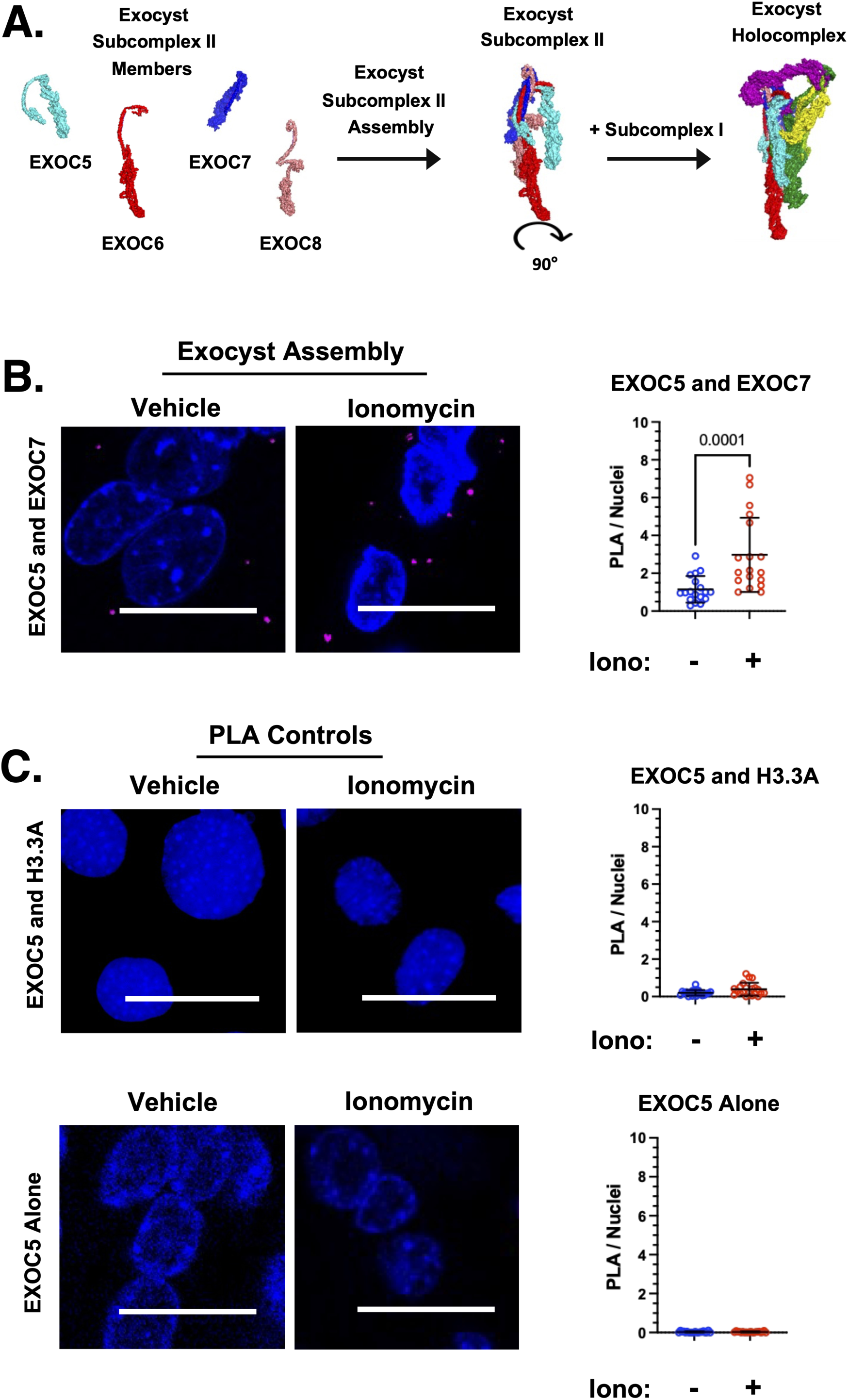
Assessing exocyst complex subunit proximity in differentiated C2C12 myotubes in response to ionomycin treatment. A. Schematic of exocyst subcomplex II members (EXOC5-8) during assembly of the 8-member holocomplex (PDB 5YFP). B. Proximity ligation assay (PLA) was used to assess EXOC5 and EXOC7 protein proximity within ∼40 nm in differentiated C2C12 myotubes following treatment with vehicle or ionomycin. Representative confocal images and quantification of PLA signal are shown. Mann–Whitney U test revealed a significant increase in EXOC5–EXOC7 proximity following ionomycin treatment (p = 0.0001). C. PLA controls included EXOC5 paired with H3.3A (histone H3) and EXOC5 antibody alone. Neither EXOC5–H3.3A proximity (p = 0.0648) nor EXOC5 antibody-alone controls (p = 0.7095) differed significantly between vehicle and ionomycin treatment. Scale bar, 25 μm; DAPI, blue; PLA puncta, red. Data were analyzed using two-tailed Mann–Whitney U tests and are shown as mean ± SD. For EXOC5–EXOC7, n = 21 fields of view per treatment; for EXOC5–H3.3A, n = 21 fields of view per treatment; and for EXOC5 antibody-alone controls, n = 21 fields of view per treatment, collected from 3 independent biological replicates.

### Rab27a exhibits a Ca^2+^ sensitive association with MVBs

The identity of Rab GTPases is intrinsically linked with specific roles in vesicular trafficking ranging from the regulation of endosomal recycling to lysosomal routing and MVB trafficking. To determine which small GTPases may participate in MVB trafficking in skeletal muscle cells, we screened for associations between Rab GTPases Rab8a, Rab11a, Rab27a, Rab27b, and Rab35 and the MVB/EV marker, CD63. We used PLA to visualize and quantify fluorescent puncta indicative of a physical proximity of less than 40 nm between CD63 and the Rab GTPases of interest (**Fig. 4A**). Our positive control for PLA included an antibody pair between CD63 and Tumor Susceptibility Gene 101 (TSG101), a member of the Endosomal Sorting Complex Required for Transport (ESCRT) EV biogenesis pathway and well-established MVB/EV marker (33). As a negative control for the Rab GTPase-CD63 PLA, we used CD63 paired with H3.3A (Histone H3) and anti-CD63 antibody alone (**Fig. 4B**). Upon quantification of PLA puncta/nuclei between vehicle and ionomycin-treated C2C12 myotubes, we found that associations between Rab27a and CD63 were significantly decreased upon ionomycin treatment (**Fig. 4A, C**). We also discovered that associations between Rab8a and CD63 significantly rose in ionomycin-treated myotubes versus vehicle control. Given the significant shift in PLA puncta/nuclei in response to Ca^2+^ flux, as well as the canonical role of Rab27a in EV biogenesis (34–36), we pursued further investigation of Rab27a-associated MVB trafficking.

**Figure 4.**
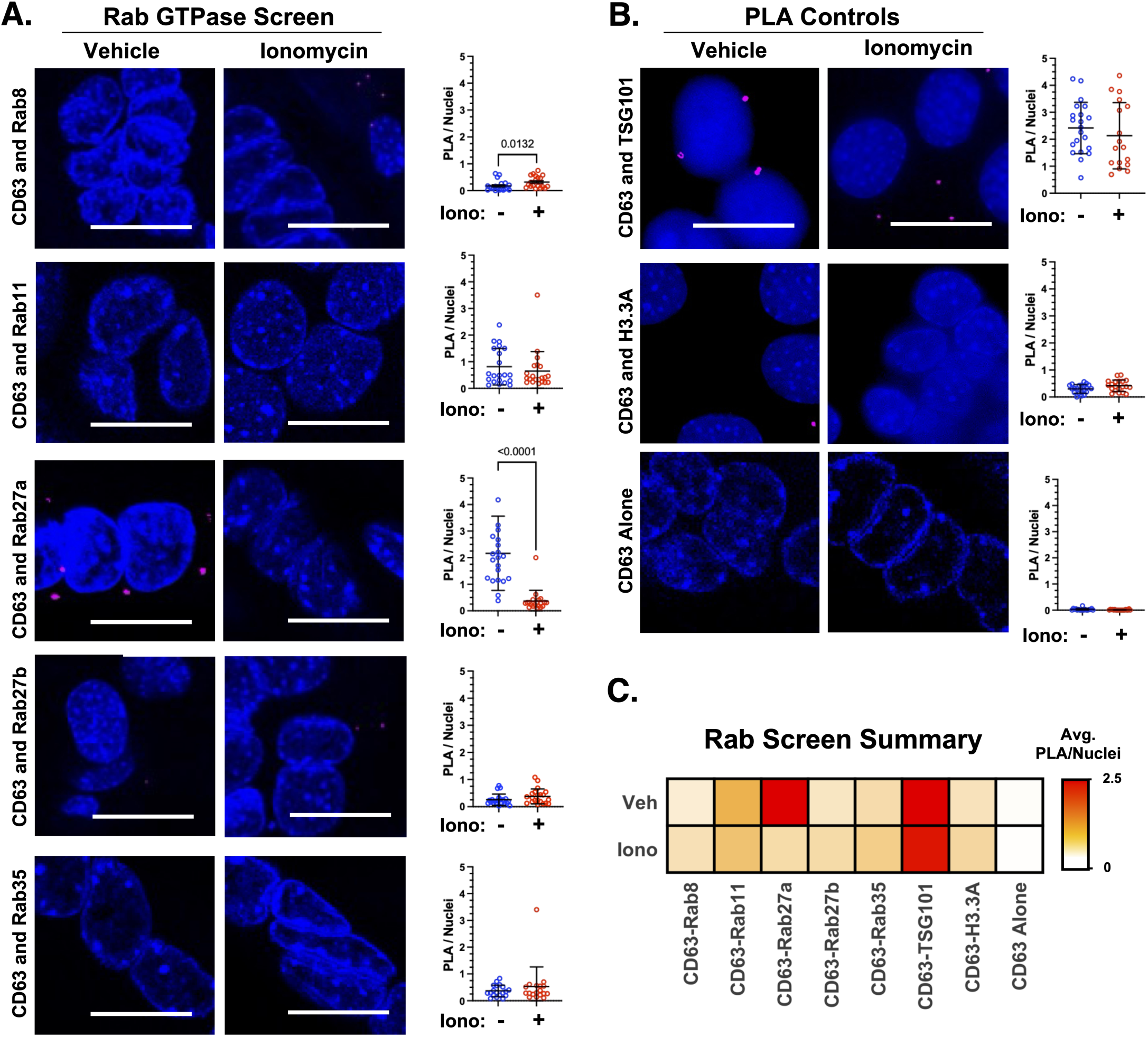
Screening for small GTPases that may direct MVB trafficking in response to Ca^2+^ signaling. A. Representative confocal images and quantification of proximity ligation assay (PLA) puncta assessing proximity between CD63 and the small GTPases Rab8, Rab11, Rab27a, Rab27b, and Rab35 in differentiated C2C12 myotubes following vehicle or ionomycin treatment. B. Representative confocal images and quantification of PLA signal for control conditions, including CD63 paired with TSG101, CD63 paired with H3.3A (histone H3), and CD63 antibody alone. C. Heatmap displaying mean PLA puncta-to-nuclei signal across conditions. Scale bar, 25 μm; DAPI, blue; PLA puncta, red. Statistical significance was assessed using two-tailed Mann–Whitney U tests, except for CD63–H3.3A, which was analyzed using an unpaired t test. Ionomycin significantly increased CD63–Rab8 (p = 0.0132) and decreased CD63–Rab27a (p < 0.0001) proximity, whereas no significant differences were observed for CD63–Rab11 (p = 0.4068), CD63–Rab27b (p = 0.1585), CD63–Rab35 (p = 0.9581), CD63–TSG101 (p = 0.3258), CD63–H3.3A (p = 0.0505), or CD63 antibody-alone controls (p = 0.1906). Data are shown as mean ± SD; sample sizes varied following outlier exclusion and are indicated by the individual analyses (n = 21 fields of view per treatment, collected from 3 independent biological replicates).

### Rab27a participates in coordinated MVB trafficking with the exocyst

Rab GTPases associate with both enzymatic and structural partners to accomplish the mobilization of vesicle-bound cargo into different cellular compartments. Expression and activity of GTPase activating proteins (GAPs), guanine exchange factors (GEFs), guanine dissociation inhibitors (GDIs), geranyl-geranyl transferase II (GGTase II) and effectors regulate the activity and localization of Rab27a (37, 38). The exocyst complex has been previously shown to interact with the Rab27a effectors melanophilin and exophilin (synaptotagmin-like protein; SYTL) in melanocytes (39) to dynamically tether melanosomes to the cell membrane. However, direct coordination between the exocyst and Rab27a in MVB trafficking has not been described, representing a gap in our mechanistic understanding of the process. To assess possible molecular coordination between the exocyst and Rab27a during MVB trafficking and tethering in response to Ca^2+^ signaling, we first used PLA to detect molecular associations between these proteins in differentiated C2C12 myotubes following ionomycin treatment. We assessed the physical proximity between Rab27a, the exocyst subunit EXOC5 and the MVB marker CD63 in a pairwise manner with or without ionomycin treatment in the presence or absence of the exocyst inhibitor ES2. Quantification of PLA puncta/nuclei between myotubes probed for EXOC5 and CD63 proximity demonstrated no significant changes between treatment with vehicle and ionomycin (**Fig. 5A**). Treatment with ES2 alone was associated with a significant rise in puncta/nuclei when compared to vehicle treatment, suggesting an increased proximity between EXOC5 and CD63 under these conditions. In addition, C2C12 myotubes co-treated with ES2 and ionomycin demonstrated a significant rise in EXOC5 and CD63 puncta/nuclei relative to treatment with ES2 alone and treatment with ionomycin alone. C2C12 myotubes probed for proximity between EXOC5 and Rab27a presented a distinct rise in PLA puncta/nuclei in ionomycin-treated myotubes versus vehicle alone (**Fig. 5B**). Cells treated with ES2 only also had significantly more puncta/nuclei, indicating an increase in EXOC5 and Rab27a associations compared to vehicle treatment. The number of EXO5C-Rab27a PLA puncta/nuclei in C2C12 myotubes co-treated with ES2 and ionomycin did not significantly differ from either ionomycin treatment or ES2 treatment. When probing for associations between CD63 and Rab27a, quantification of PLA puncta revealed no significant changes between treatments with vehicle, ionomycin or ES2 alone (**Fig. 5C)**. While we observed a similar ionomycin-induced reduction of CD63-Rab27a proximity to our original Rab screen, this difference did not reach statistical significance when analyzed within the four-condition experimental design. Of note, ES2 treatment paired with ionomycin resulted in a significant rise in the number of puncta/nuclei between CD63 and Rab27a relative to treatment with either ES2 alone or ionomycin alone, demonstrating an increase in proximity between the GTPase and the CD63-labeled MVBs. Our results indicate that MVBs are spatially coupled between Rab27a and the exocyst in a manner that is dependent on Ca^2+^ signaling and exocyst activity.

**Figure 5.**
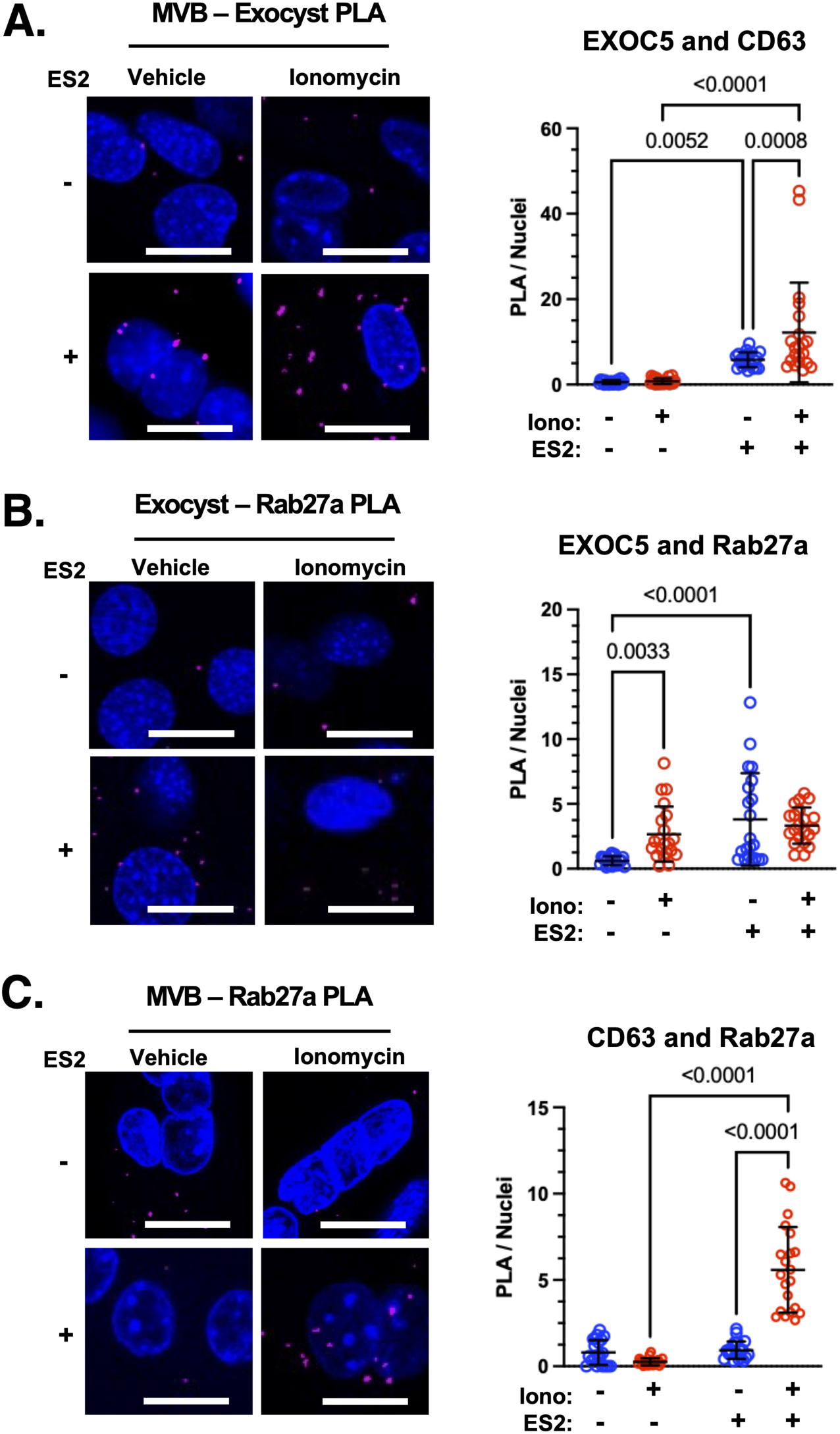
Evaluating the molecular associations between the GTPase Rab27a, the exocyst subunit EXOC5, and the MVB marker CD63. A-C. Representative confocal images and quantification of proximity ligation assay (PLA) signal assessing proximity between EXOC5 and CD63 (A.), EXOC5 and Rab27a (B.), and CD63 and Rab27a (C.) in differentiated C2C12 myotubes following treatment with vehicle (EtOH) or ionomycin in the presence of DMSO or Endosidin 2 (ES2). Scale bar, 50 μm; DAPI, blue; PLA puncta, red. For EXOC5-CD63 proximity (A), two-way ANOVA revealed significant effects of ES2 treatment (F(1,80) = 41.64, p < 0.0001), ionomycin treatment (F(1,80) = 6.540, p = 0.0124), and the ES2 × ionomycin interaction (F(1,80) = 5.722, p = 0.0191), followed by uncorrected Fisher’s LSD multiple-comparisons tests. Ionomycin significantly increased EXOC5–CD63 proximity in the presence of ES2 (p = 0.0008) but not DMSO (p = 0.9073). ES2 significantly increased EXOC5–CD63 proximity under both vehicle (p = 0.0052) and ionomycin (p < 0.0001) conditions. For EXOC5–Rab27a proximity (B), two-way ANOVA revealed a significant effect of ES2 treatment (F(1,80) = 16.11, p = 0.0001) and a significant ES2 × ionomycin interaction (F(1,80) = 6.943, p = 0.0101), but no significant main effect of ionomycin treatment (F(1,80) = 2.706, p = 0.1039). Ionomycin significantly increased EXOC5–Rab27a proximity in the presence of DMSO (p = 0.0033) but not ES2 (p = 0.4860). ES2 significantly increased EXOC5–Rab27a proximity under vehicle conditions (p < 0.0001) but not ionomycin conditions (p = 0.3324). For CD63–Rab27a proximity (C), two-way ANOVA revealed significant effects of ES2 treatment (F(1,80) = 89.85, p < 0.0001), ionomycin treatment (F(1,80) = 51.11, p < 0.0001), and the ES2 × ionomycin interaction (F(1,80) = 81.31, p < 0.0001). Ionomycin did not significantly alter CD63–Rab27a proximity in the presence of DMSO (p = 0.1904) but significantly increased proximity in the presence of ES2 (p < 0.0001). ES2 did not significantly alter CD63–Rab27a proximity under vehicle conditions (p = 0.7449) but significantly increased proximity following ionomycin treatment (p < 0.0001). Data represent n = 3 independent biological replicates, with 7 fields of view quantified per replicate (n = 21 fields of view per condition), and are shown as mean ± SD.

### Immobilized protein binding assay supports direct Rab27a-EXOC6 interaction

Given that the PLA data suggest a molecular association between Rab27a and exocyst complex members, we also performed a co-immunoprecipitation (CoIP) to attempt to recover binding partners when targeting exocyst subunits. We utilized a well-validated antibody against EXOC4 to immunoprecipitate endogenous EXOC4 and associated proteins from wild-type (WT) C2C12 myotube lysates. Beads-only and isotype IgG controls were included to assess nonspecific protein recovery attributable to the beads or immunoglobulin, respectively (**Fig. 6A**). When probing for EXOC4, we observed a band in all three input lanes and in our EXOC4 IP lane, confirming the presence of this subunit in the lysates and the successful immunoprecipitation of this exocyst subunit. Some non-specific fluorescent signal in all three IP lanes may be due to denatured, reactive Fc molecules derived from the immuno-capture beads. When probing for Rab27a, we observed bands in all three lysate input lanes and additionally in our EXOC4 antibody IP lane. Although Rab27a co-precipitated with exocyst subunit EXOC4, this interaction may be indirectly modulated by other proteins, including other exocyst subunits. To assess the potential of a direct interaction between Rab27a and members of the exocyst complex, we tested if direct binding was achievable between Rab27a and isolated components of the exocyst complex. We selected EXOC6 as a binding substrate for our direct binding assays based on its established role as an effector of multiple Rab GTPases. We immobilized purified EXOC6-FlagTag and titrated in recombinant BSA, 6xHis-Rab27a, 6xHis-GFP, or GST-EXOC5 to test if a combination with EXOC6 would produce a binding isotherm. We found that there was a dose-dependent rise in fluorescent signal corresponding to both Rab27a and EXOC5 titration, suggesting a concentration-dependent complex formation between EXOC6 and Rab27a in addition to EXOC6 and EXOC5, our positive control (**Fig. 6B**). Comparison of nucleotide-loaded Rab27a further revealed distinct binding profiles between GTPγS- and GDPβS-loaded Rab27a, suggesting that the interaction with EXOC6 may be influenced by the nucleotide state of Rab27a. Our data also demonstrates that there was no concentration-dependent change in signal in titrations of BSA or 6xHis-GFP negative controls.

**Figure 6.**
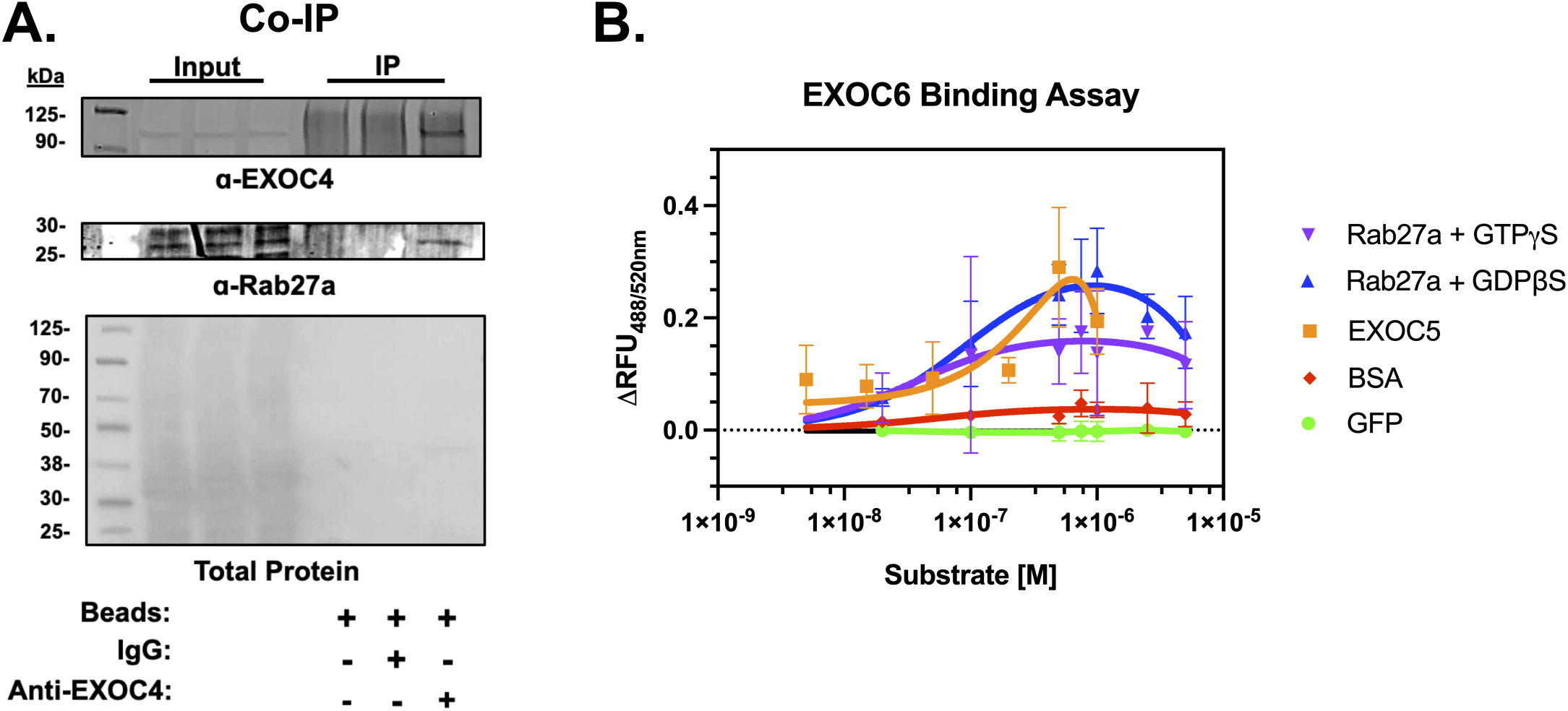
Binding characterization of EXOC6 with BSA, Rab27a, GFP and EXOC5. A. Western analysis of proteins immunoprecipitated from differentiated C2C12 myotubes: in vitro interactions were tested by immunoprecipitation of EXOC4 using an anti-EXOC4 antibody, with isotype control IgG and beads only as the negative controls. Immunoprecipitates were subjected to immunoblot analysis using antibodies against EXOC4 and Rab27a. B. Using a solid-phase binding assay, we tested if the purified exocyst subunit EXOC6 would produce concentration-dependent binding with purified BSA, Rab27a, GFP and EXOC5. Data were fit to a Langmuir Isotherm model to produce a binding curve. Data shown are averages of 3 independent experiments, each performed with technical triplicates (n = 3). Rab27a GTPγS: Purple, Rab27a GDPβS: Blue, BSA: Red, Rab27a: Blue, GFP: Green, EXOC5: Orange. Data are shown as mean ± SD.

### Molecular modeling supports a direct interaction between Rab27a and exocyst subunits

To interrogate the atomic-scale coordination between Rab27a and the exocyst complex, we utilized molecular modeling to estimate the dissociation constant (predicted K_D_) and residue-specific coordination (within 10Å) between the exocyst complex and Rab27a. We separately paired the eukaryotic structure of the exocyst complex (Protein Data Bank (PDB) 5YFP (18)) against the Fc domain of human immunoglobulin G (IgG) (PDB 4QGT (40)) and human Rab27a (modeled via AlphaFold3 (41)) (**Fig. 7A**). We compared estimated dissociation constants between the exocyst-IgG and exocyst-Rab27a pairs and found that the exocyst-Rab27a estimated K_D_ values ((1.75 ± 2.03) × 10^-6^ M) was significantly lower than the estimated K_D_ values from the exocyst-IgG complex ((3.42 ± 4.15) × 10^-5^ M) (**Fig. 7B**). We also modeled complexes between the exocyst and other Rab GTPases including Rab8, Rab10 and Rab11 that are known to use the exocyst complex as their effector (15, 22, 39, 42–44). We dissected the 3D coordinates from our modeling efforts between the Rab GTPases and the exocyst and found that there are shared regions of binding between the Rab GTPases and exocyst subunits EXOC1, 2, 5 and 8 (**Fig. 7C**). Lastly, we determined the estimated K_D_ (M) for the exocyst-Rab GTPase interactions and found no significant differences in predicted strength of binding observed between the models (**Fig. 7D**) (Rab8: (3.50 ± 4.91) × 10^-6^ M, Rab10: (2.29 ± 2.88) × 10^-6^ M, Rab11: (1.21 ± 1.08) × 10^-6^ M). Together, we found homologous binding topology and conserved binding energetics between the exocyst and Rab GTPase complexes that supports the direct binding between Rab27a and the exocyst subunits.

**Figure 7.**
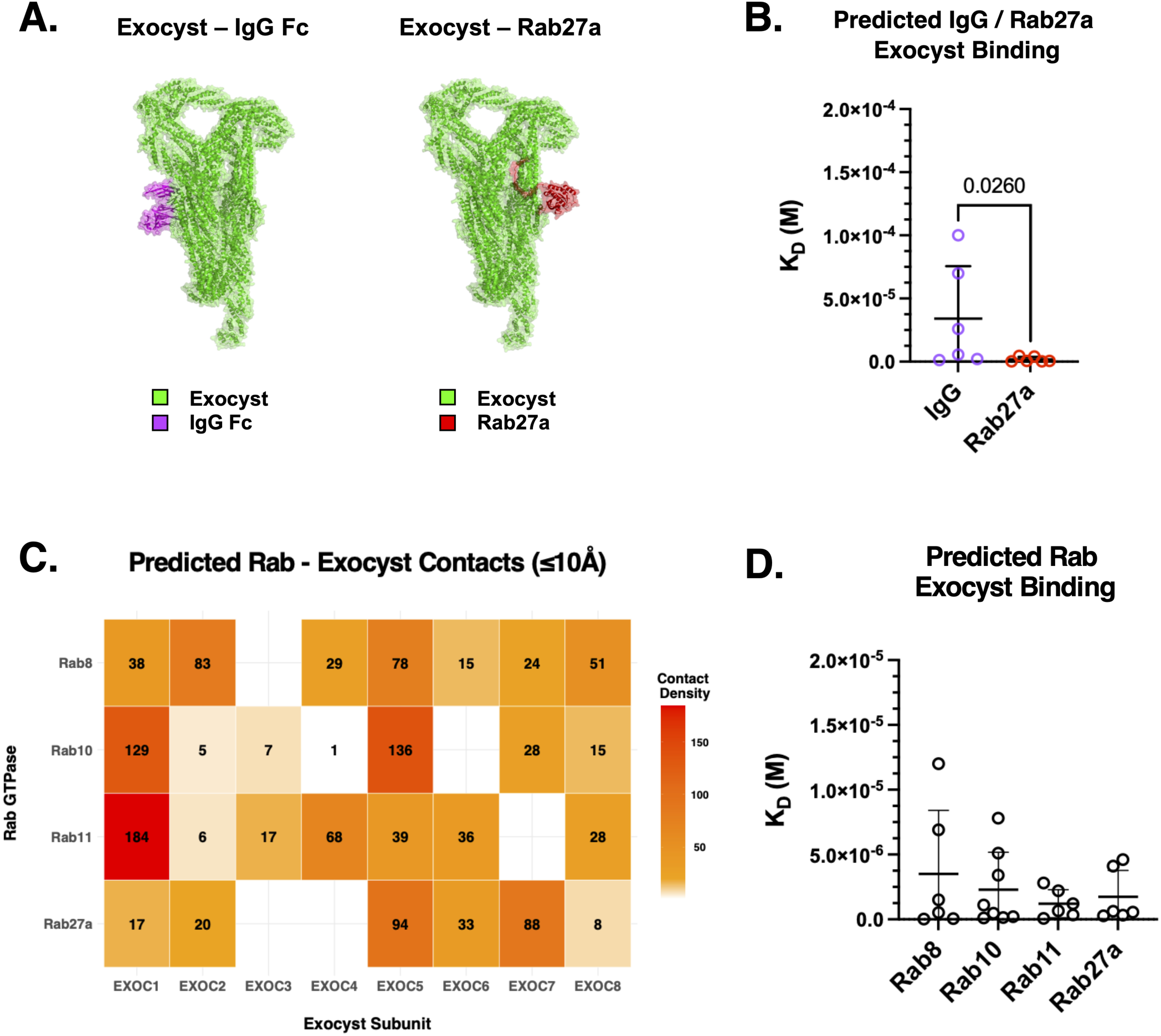
Molecular modeling of the interaction between Rab27a and exocyst subunits. A. Molecular models of the exocyst holocomplex (PDB 5YFP) in association with either the IgG Fc domain (PDB 4QGT) as a control or Rab27a modeled using AlphaFold3. B. Comparison of estimated dissociation constants (K_D_) for exocyst complex formation with IgG Fc or Rab27a. Estimated K_D_ values were significantly lower for the predicted Rab27a–exocyst complexes compared with the IgG Fc–exocyst control (two-tailed Mann–Whitney U test, p = 0.0260). n = 6 independent computational solutions/poses per condition. C. Heatmap depicting estimated K_D_ values for interactions between individual exocyst subunits (EXOC1–EXOC8) and the Rab GTPases Rab8, Rab10, Rab11, and Rab27a. D. Comparison of estimated K_D_ values for predicted exocyst–Rab complex formation across Rab8, Rab10, Rab11, and Rab27a. A Kruskal–Wallis test revealed no significant difference among the four Rab GTPases (p = 0.9668), and Dunn’s multiple-comparisons tests revealed no significant pairwise differences (all adjusted p > 0.9999). n = 6-8 independent computational solutions/poses per Rab GTPase. Data are shown as mean ± SD.

### Evaluating the exocyst complex as a Ca^2+^-dependent Rab27a effector

While the solid-phase binding assay and molecular modeling approaches provided complementary quantitative and structural insights into the association between Rab27a and the exocyst, we also sought to understand how Ca^2+^ signaling could potentially reshape the proximity networks of Rab27a and the exocyst in living cells. We designed and developed several plasmids encoding a piggyBac transposase suitable for stable mammalian genomic integration (45, 46), an antibiotic (Hygromycin) resistance gene for selection and an UltraID (evolved BioID biotin ligase)-fusion protein of either Rab27a or EXOC4 with an N-terminal Myc tag epitope (**Fig. 8A**). UltraID is capable of biotinylating proteins within enzymatic proximity and possesses several distinct advantages over other proximity-labeling approaches including a small < 25kDa size, relatively fast biotinylation kinetics, a high signal-to-noise ratio, and ≤ 10 nm labeling range (47, 48). Furthermore, the strategy of using the UltraID as a fusion partner of Rab27a and EXOC4 allowed us to explore the proximity-labeled proteome of these proteins in living SkM cells as opposed to a post-lysis capture. To interrogate the role of nucleotide-binding state to Rab27a’s proximity network, we also performed mutagenesis to create Rab27a^T23N^ and impair Rab27a’s ability to bind to GTP (49). We cloned and stably expressed UltraID fusion proteins in C2C12 SkM myotubes, including EXOC4-UltraID, UltraID-Rab27a^WT^, UltraID-Rab27a^T23N^, and UltraID alone as a labeling control (**Fig. 8A**). Before establishing the labeling potential of our UltraID-Rab27a system on known Rab27a interactors, we assessed the expression and the kinetics of the biotin ligase activity of our UltraID constructs (**Supp S3**). We confirmed the presence of a-Myc tag band (N-terminus of UltraID) in transgenic cell lysates. In cases where the Myc band was not detected in transgenic cell lysate, we isolated UltraID fusion proteins by incubating lysates with NeutrAvidin beads. NeutrAvidin pulldown purified UltraID fusion proteins due to a self-biotinylation phenomenon of the UltraID-fusion protein. We also incubated UltraID-expressing cells with 50 μM biotin from 0-24 hours to assess the kinetics of labeling. We demonstrated that increased incubation with biotin resulted in an increase in fluorescent signal (**Supp S3**). After verifying the expression and biotin-responsive labeling of the UltraID fusion proteins, we performed a labeling experiment to assess the fidelity of the biotin labeling pattern and evaluate the pull-down of known interactors of Rab27a while avoiding the labeling of control proteins that are unlikely to associate with Rab27a. We performed the NeutrAvidin-mediated pull-down in the following experimental groups a.) WT C2C12 myotubes treated with biotin b.) UltraID-Rab27a^WT^-expressing myotubes treated with vehicle, and c.) UltraID-Rab27a^WT^-expressing myotubes treated with biotin (**Fig. 8B**). Despite poor baseline expression in C2C12 myotubes, NeutrAvidin capture of biotinylated proteins revealed the successful pulldown of SYTL4 (SLP4A; synaptotagmin-like family member 4), a known interactor of Rab27a (50, 51), confirming our model system can help identify interacting partners via proximity labeling. As a negative control, we also probed for GAPDH (Glyceraldehyde 3-phosphate dehydrogenase) and found it was not resolved in any of the pulldown lanes. Lastly, to follow up on our PLA results, we probed for EXOC4 and found that it was pulled down from cells expressing UltraID-Rab27a subjected to biotin treatment. Next, we set out to characterize the proximity-labeled proteomes of EXOC4-UltraID, UltraID-Rab27a, and UltraID-Rab27a^T23N^ C2C12 myotubes upon treatment with vehicle or ionomycin using data-independent acquisition (DIA) proteomics (**Fig. 8C, Supp. S4)**. We found that several exocyst subunits were significantly enriched among the proteins labeled by EXOC4-UltraID relative to UltraID alone during vehicle treatment (EXOC1, EXOC2, EXOC3, EXOC6A, EXOC6B, EXOC7 and EXOC8), further validating our proximity labeling approach. (**Fig. 8D**). We also observed that EXOC4-UltraID labeled partners of the exocyst complex, including Actin Related Protein 2/3 complex subunit 2 (ARPC2) (52), Cyclin Dependent Kinase 1 (CDK1) (19) and Tropomyosin 3 (TPM3) (53). Under these conditions, Rab27a was not significantly more abundant among the proteins labeled by EXOC4-UltraID versus UltraID alone control. During treatment with ionomycin, we observed a shift whereby the abundance of biotinylated EXOC3, EXOC6B and EXOC7 was significantly decreased compared to control conditions. Analysis of proteins significantly enriched in UltraID-Rab27a relative to UltraID alone under vehicle conditions identified several Rab27a effectors, including SYTL1, 2 and 5 as well as the exocyst complex subunit, EXOC6B (**Fig. 8E**). Ionomycin treatment was associated with a loss of significantly higher labeling of STYL1 and EXOC6B, but resulted in the significantly higher abundance of the labeled EXOC6B isoform, EXOC6. We then examined Ca^2+^-dependent differences in protein labeling by UltraID-Rab27a^WT^ and UltraID-Rab27a^T23N^ (the GDP-locked conformation of Rab27a (54, 55)) by comparing the abundance of biotinylated proteins in samples treated with vehicle or ionomycin (**Fig. 8F**). Comparison between UltraID-Rab27a^WT^ and UltraID-Rab27a^T23N^ proximitomes during vehicle treatment revealed no significant differences in the proximity of UltraID-Rab27a to exocyst subunits. However, during Ca^2+^ stimulation with ionomycin treatment, components of exocyst subcomplex I and II (EXOC1 and EXOC6) were significantly enriched in the UltraID-Rab27a^WT^-labeled samples compared to those biotinylated by UltraID-Rab27a^T23N^. We also found significant enrichment of Rab27a effectors, including SYTL1 and SYTL5, in cells expressing UltraID-Rab27a^WT^ over UltraID-Rab27a^T23N^. The significant abundance of exocyst subunits biotinylated by UltraID-Rab27a^WT^ over UltraID alone, as well UltraID-Rab27a^T23N^, further demonstrates a reasonable proximity between the active Rab27a and the exocyst in SkM cells.

**Figure 8.**
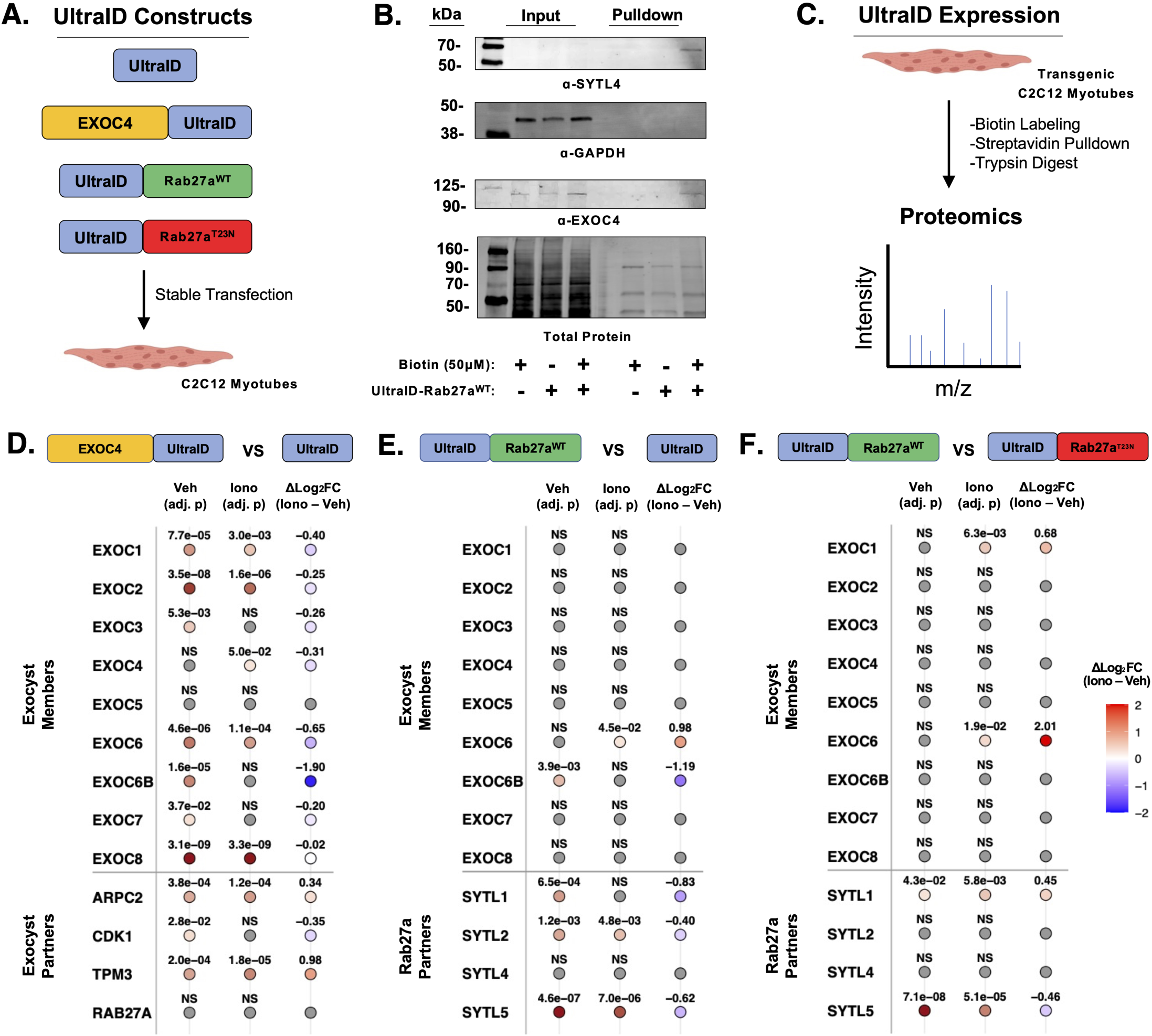
Using a biotin-labeling approach to interrogate the Ca2+-signaling-dependent proximitome of the exocyst subunit EXOC4 and Rab27a. A. Schematic of UltraID fusion proteins including EXOC4-UltraID, UltraID-Rab27aWT, UltraID-Rab27aT23N and UltraID alone expressed in C2C12 myotubes. B. Western blot of UltraID-Rab27a pulldowns from WT and UltraID-Rab27aWT-expressing C2C12 myotubes treated ± biotin (50 μm); membranes were probed for SYTL4 (Synaptotagmin-Like Protein-4), GAPDH, and EXOC4. C. Experimental outline of UltraID-based proximity labeling and data-independent acquisition (DIA) proteomic analysis. D-F. Plots showing normalized biotinylated proteins enriched through proximity labeling by EXOC4-UltraID relative to UltraID alone during vehicle and ionomycin treatment (D), UltraID-Rab27aWT vs UltraID alone (E), and UltraID-Rab27aWT vs UltraID-Rab27aT23N (F). Shaded dots represent proteins significantly labeled vs UltraID (P adj < 0.05). Categories for proteins include exocyst members [EXOC1-8] and exocyst partners [Actin Related Protein 2/3 complex subunit 2 (ARPC2), Cyclin Dependent Kinase 1 (CDK1) and Tropomyosin 3 (TPM3)]. Rab27a partners include SYTL1, 2, 4 and 5. Significant p adj. < 0.05 Data represent n = 4 independent biological experiments. Data are shown as mean ± SD.

### Real-time measurements of Ca^2+^-regulated CD63 exocytosis using a fluorescent reporter

To better understand the real-time dynamics of membrane fusion and EV secretion events, we cloned a CD63 transgene from a lentiviral delivery plasmid(56) that expresses a pH-sensitive GFP (pHluorin) within the first extracellular loop of CD63 and a pH-stable RFP (mScarlet) on CD63’s C-terminus. We created a vector featuring a piggyBac transposase, an antibiotic resistance gene and the pHluorin-CD63-mScarlet for stable transfection of our SkM cell line (**Supp. S5A**). The advantage of this construct resides in the pH-dependent fluorescent activity of the pHluorin fluorescent molecule appended to CD63’s first extracellular domain that is quenched in the acidic environments of endosomal and MVB-compartments. Upon fusion between the plasma membrane and the MVB membrane, the CD63 molecule’s N-terminus on the surface of the small EV encounters a pH shift, which allows for an increase in pHluorin’s fluorescence intensity (**Fig. 9A**) (56–58). To establish a stable, pHluorin-CD63-mScarlet expressing C2C12 cell line, we transfected C2C12 myoblasts with the pHluorin-CD63-mScarlet plasmid and then used hygromycin treatment as an antibiotic selection pressure to create a stable polyclonal fluorescent line. Given the importance of homogeneity in fluorescence-based biochemical experiments, we used flow cytometry-assisted cell sorting (FACS) to sort for pHluorin-CD63-mScarlet expressing, viable single cells to generate a monoclonal pHluorin-CD63-mScarlet C2C12 cell line (**Supp. S5B**). We validated transgene expression in our monoclonal cells and confirmed that dual-tagged CD63 is enriched in small EVs isolated from pHluorin-CD63-mScarlet C2C12 cells **(Supp. S5C-E)**. We then tested if we could induce changes in pHluorin-CD63 fluorescence (ΔpHluorin-CD63) that reflect MVB fusion and EV secretion events during ionomycin treatment. Importantly, our monoclonal pHluorin-CD63-mScarlet C2C12 myotubes responded to incubations with a range of ionomycin concentrations (0, 1, 5, and 10 μM) by producing a semi-dose responsive change in pHluorin-CD63 (ΔpHluorin-CD63) fluorescence (**Fig. 9B**). We then measured ΔpHluorin-CD63 fluorescence in response to ionomycin treatment in the presence of known inhibitors of EV secretion. We pre-treated pHluorin-CD63-mScarlet C2C12 myotubes with either vehicle, 10 μM Dantrolene (the small molecule inhibitor of the Ryanodine Receptor (59)) or 10 μM Manumycin A (an inhibitor of the Ras signaling cascade utilized during EV biogenesis (60)). We found that ΔpHluorin-CD63 fluorescence was significantly stymied in response to ionomycin after 15 minutes of treatment only in C2C12 cells co-treated with Manumycin A. The effects of ionomycin were not significantly altered in Dantrolene-treated cells (**Fig. 9C**). We additionally included an experiment again testing the effects of exocyst inhibition with ES2 and found that co-treatment of ES2 and ionomycin was associated with a significantly lower ΔpHluorin-CD63 when compared to treatment with ionomycin alone (**Fig. 9D**), supporting our previous findings on the critical role of exocyst activity during EV release in response to ionomycin treatment in differentiated pHluorin-CD63-mScarlet C2C12 myotubes.

**Figure 9.**
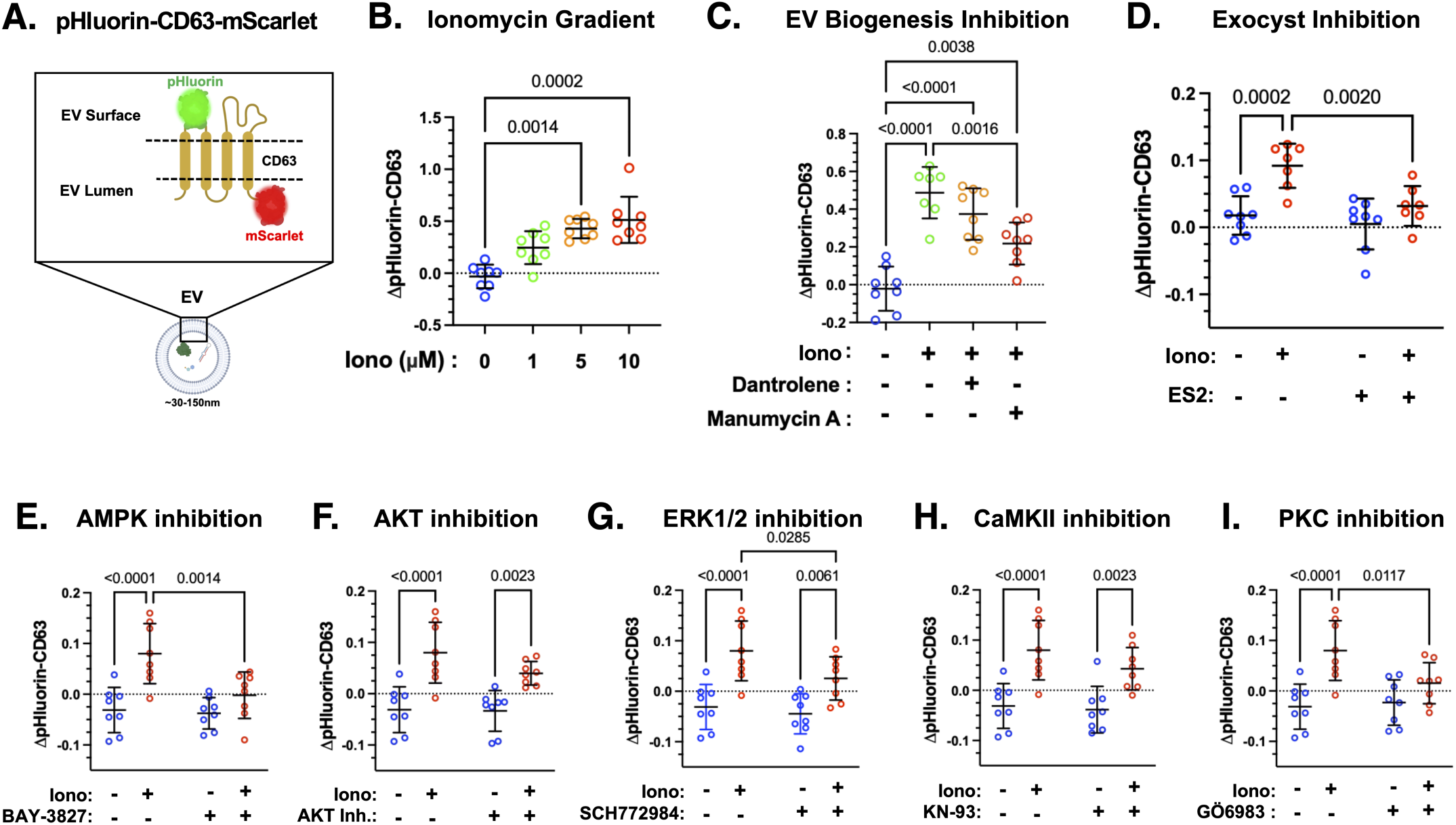
Using a fluorescent reporter cell line to identify the signaling pathways involved in Ca^2+^ flux-induced EV release in muscle cells. A. Schematic representation of the dual-tagged EV reporter pHluorin-CD63-mScarlet and its orientation relative to the EV membrane. B. Quantification of changes in pHluorin fluorescence (ΔpHluorin-CD63; excitation, 488 nm; emission, 511 nm; arbitrary fluorescence units [AFU]) following treatment with 0–10 μM ionomycin for 15 min. Kruskal–Wallis analysis revealed a significant difference among treatment groups (p = 0.0001), followed by Dunn’s multiple-comparisons tests. Ionomycin significantly increased ΔpHluorin-CD63 at 5 μM (p = 0.0014) and 10 μM (p = 0.0002) relative to 0 μM ionomycin, with no significant difference between 5 and 10 μM ionomycin (p > 0.9999). C. ΔpHluorin-CD63 following ionomycin treatment with or without pretreatment with the Ca²⁺ release inhibitor dantrolene or the Ras inhibitor Manumycin A. One-way ANOVA revealed a significant difference among treatment groups (F(3,27) = 23.53, p < 0.0001), followed by Tukey’s multiple-comparisons test. Ionomycin significantly increased ΔpHluorin-CD63 relative to untreated controls (p < 0.0001). Dantrolene did not significantly alter the ionomycin-induced response (p = 0.3234), whereas Manumycin A significantly reduced the response relative to ionomycin alone (p = 0.0016). D. ΔpHluorin-CD63 following ionomycin treatment in the presence of DMSO or Endosidin 2 (ES2). Two-way ANOVA revealed significant main effects of ES2 treatment (F(1,26) = 9.308, p = 0.0052) and ionomycin treatment (F(1,26) = 17.95, p = 0.0003), but no significant ES2 × ionomycin interaction (F(1,26) = 3.913, p = 0.0586), followed by uncorrected Fisher’s LSD multiple-comparisons tests. Ionomycin significantly increased ΔpHluorin-CD63 in DMSO-treated cells (p = 0.0002) but not ES2-treated cells (p = 0.1222). ES2 significantly reduced ΔpHluorin-CD63 under ionomycin conditions (p = 0.0020) but did not significantly alter the response under vehicle conditions (p = 0.4394). E–I. ΔpHluorin-CD63 following ionomycin treatment in the presence or absence of small-molecule inhibitors targeting AMPK (BAY-3827; E), AKT (AKT Inhibitor VIII; F), ERK1/2 (SCH772984; G), CaMKII (KN-93; H), or PKC (Gö6983; I). Two-way ANOVA followed by uncorrected Fisher’s LSD multiple-comparisons tests was used for panels E–I. E. For AMPK inhibition, a significant AMPK inhibitor × ionomycin interaction was observed (F(1,28) = 5.313, p = 0.0288). Ionomycin significantly increased ΔpHluorin-CD63 in DMSO-treated cells (p < 0.0001) but not in BAY-3827-treated cells (p = 0.1333), and BAY-3827 significantly reduced the ionomycin-associated response (p = 0.0014). F. For AKT inhibition, a significant AKT inhibitor × ionomycin interaction was observed (F(1,28) = 19.95, p = 0.0001). Ionomycin increased ΔpHluorin-CD63 in both DMSO-(p = 0.0064) and AKT Inhibitor VIII-treated cells (p < 0.0001), with a significantly greater response following AKT inhibition under ionomycin conditions (p < 0.0001). G. For ERK1/2 inhibition, ionomycin significantly increased ΔpHluorin-CD63 in both DMSO-(p < 0.0001) and SCH772984-treated cells (p = 0.0061). SCH772984 significantly reduced the ionomycin-treated signal relative to DMSO (p = 0.0285), although the inhibitor × ionomycin interaction was not significant (F(1,28) = 1.509, p = 0.2295). H. For CaMKII inhibition, ionomycin significantly increased ΔpHluorin-CD63 in both DMSO-(p < 0.0001) and KN-93-treated cells (p = 0.0023), with no significant effect of KN-93 under either vehicle (p = 0.7648) or ionomycin (p = 0.1387) conditions; the inhibitor × ionomycin interaction was not significant (F(1,28) = 0.7465, p = 0.3949). I. For PKC inhibition, a significant PKC inhibitor × ionomycin interaction was observed (F(1,28) = 4.598, p = 0.0408), with a significant main effect of ionomycin treatment (F(1,28) = 19.49, p = 0.0001) but no significant main effect of Gö6983 treatment (F(1,28) = 2.785, p = 0.1063). Ionomycin significantly increased ΔpHluorin-CD63 in DMSO-treated cells (p < 0.0001) but not in Gö6983-treated cells (p = 0.1196). Gö6983 significantly reduced the ΔpHluorin-CD63 response under ionomycin conditions (p = 0.0117) but did not significantly alter the response under vehicle conditions (p = 0.7393). Data are shown as mean ± SD. Each panel represents one representative experiment performed with 8 technical replicate wells per condition; 3 independent experiments were performed in total.

We then revisited the kinases AMP-activated protein kinase (AMPK), protein kinase B (AKT), extracellular signal-regulated kinase 1/2 (ERK1/2), who’s altered phosphorylation status we detected before in ionomycin treated C2C12 myotubes and assessed their contribution to EV release using the pHluorin-CD63-mScarlet C2C12 myotubes. Additionally, we tested calcium/calmodulin-dependent protein kinase II (CaMKII) and protein kinase C (PKC), which have direct roles in regulating Ca^2+^-dependent signaling and organization of secretion and contraction machinery. We challenged the ΔpHluorin-CD63 fluorescence produced by ionomycin treatment with kinase inhibitors against AMPK (BAY-3827 (61)), AKT (AKT Inhibitor VIII (62)), ERK1/2 (SCH772984 (63)), CaMKII (KN-93 (64)) and PKC (Gö-6983 (65)). Treatment of myotubes with any of the inhibitors alone did not produce significant changes in pHluorin-CD63 fluorescence when compared to vehicle. We found that pharmacological inhibition of AMPK, ERK1/2 and PKC with ionomycin co-treatment was associated with a significant impairment of the ΔpHluorin-CD63 elicited by ionomycin alone (**Fig. 9E**), but inhibition of AKT or CaMKII did not result in a significant decrease in ΔpHluorin-CD63 fluorescence in response to ionomycin treatment after 15 minutes. With our EV secretion reporter cell line, we observed distinct profiles of ΔpHluorin-CD63 fluorescence associated with chemical inhibition of key kinases such as AMPK, ERK1/2, and PKC that relate to Ca^2+^ regulation and cytoskeletal organization.

## DISCUSSION

Our study identifies a Ca^2+^-responsive trafficking network that coordinates the secretion of small (∼50-150 nm) EVs in SkM cells. Although the introduction of Ca^2+^ has been demonstrated to stimulate EV secretion, the trafficking machinery linking Ca^2+^-signaling to MVB trafficking and subsequent EV secretion remains incompletely understood. Here, we demonstrate that Ca^2+^ signaling promotes EV secretion through dynamic regulation of Rab27a-exocyst interactions.

First, we show that treatment with the ionophore, ionomycin, is tied to the activation of kinases and GTPase activating proteins (GAPs) prominently featured in Ca^2+^-dependent pathways such as AMPK (66–69), ERK1/2 (67, 68), and AS160 (69) (**Fig. 1C-F**) in cultured C2C12 myotubes. We observed significantly higher levels of AMPK phosphorylation on Thr172, typically phosphorylated upon AMP sensing via LKB1 and Ca^2+^ sensing via CaMKKβ (Ca^2+^/Calmodulin-dependent protein kinase beta chain (70)). Consistent with the engagement of Ca^2+^-responsive kinase pathways, our observation of increased phosphorylation of ERK1/2 at Thr202 and Tyr204 following ionomycin treatment may be explained by activation by Ca^2+^/calmodulin-dependent signaling pathways, including CaMKKβ-dependent mechanisms that act on the RAS-RAF-MEK signaling cascade (71, 72). In ionomycin-treated myotubes, our observation of increased AS160 phosphorylation points to the role of Thr642 as a regulatory site on AS160 whose phosphorylation inhibits AS160’s ability to sequester Rab GTPases that are recruited to traffic vesicles (73). Previous studies have also reported that elevated intracellular Ca^2+^ can reduce AKT Ser473 phosphorylation, similar to our observation, potentially through altered phosphoinositide interactions that impair AKT recruitment to the plasma membrane (74). The competitive occupation of AKT’s PIP_3_ binding domain can affect proper localization of AKT to the plasma membrane, which may reduce access to kinases responsible for AKT phosphorylation. In addition to biophysical restriction of AKT mobility, enzymatic regulators can also limit AKT activation. Alternatively, activation of Ca^2+^-responsive phosphatases such as calcineurin may contribute to the reduced AKT phosphorylation observed following ionomycin treatment (75, 76). Although AS160 is a phospho-substrate of AKT, AMPK activity has also been shown to mediate AS160 phosphorylation (77), and this could possibly occur during intense rises in intracellular Ca^2+^ and in the presence of activated AMPK. Furthermore, phosphorylation of AS160 is generally associated with enhanced Rab-dependent vesicle trafficking (78). Through treatment of C2C12 myotubes with the Ca^2+^ ionophore, ionomycin, we are able to confirm dynamic regulation of kinases and GTPase-activating proteins essential to contraction-related signaling pathways.

Second, we found that treatment with ionomycin, in addition to activating signal transduction pathways in cultured SkM cells, also leads to a significant increase in the number of small EVs released (**Fig. 2C**). Our detection of these EVs offers an exclusive window into the ∼50-150 nm size vesicle population as our EV isolation methodology via size-exclusion chromatography enriches particles within this size range. We are experimentally agnostic to the potential secretion dynamics of other classes of particles that exceed or fall below our selected size regime, as those remained uncaptured and unmeasured. The increase in EV size observed following ionomycin treatment may reflect Ca^2+^-sensing proteins that respond to membrane damage and influence membrane structure. Annexins (ANXA4-6), for instance, have been implicated in membrane fusion, membrane curvature, and cytoskeletal remodeling associated with EV biogenesis and release (79–81). Elevated intracellular Ca^2+^, therefore, may be accompanied by significant membrane reorganization and may account for changes in EV size.

Third, we identified that Ca^2+^ signaling promotes exocyst assembly, and exocyst function is required for the efficient secretion of small EVs in response to ionomycin-induced Ca^2+^ signaling. Inhibition of the exocyst complex via ES2 treatment leads to a significant decrease in small EV secretion, even during co-treatment with ionomycin. Our pharmacological inhibition of the exocyst reflects the importance of exocyst function in EV secretion, as evidenced by earlier reports of a consequential decrease in EV secretion following knockdown of exocyst subunits in various cell models (16, 28). This hypothesis also aligns with our observation that the intra-exocyst subunit proximity between EXOC5 and EXOC7 after treatment with ionomycin may reflect a Ca^2+^-sensitive rearrangement of exocyst subunits in a way that increases the spatial proximity of subcomplex II members EXOC5 and EXOC7 (**Fig. 3B**). In addition to belonging to the same subcomplex, subunits EXOC5 and EXOC7 represent separate but critical aspects of exocyst function. EXOC5 expression has been demonstrated to be required for exocyst complex stability, as downregulation of EXOC5 (via RNAi) leads to the downregulation of other exocyst subunits (27, 82, 83). EXOC7 binds to lipid moieties, such as phosphoinositides, thereby allowing plasma membrane tethering and polarized trafficking of vesicular cargo (16, 84). We offer two distinct and possibly co-occurring explanations for the phenomenon of increased subcomplex proximity during Ca^2+^ signaling; modulation of exocyst subunit proximity through direct signaling-dependent and membrane-dependent mechanisms. Firstly, exocyst complex assembly can be regulated via direct phosphorylation of exocyst subunits. ERK1/2 and PKC (Protein Kinase C), which both respond to increases in Ca^2+^, can phosphorylate exocyst subunits (EXOC2, EXOC7) and allow for the tuning of complex assembly (20, 21). Assembly of subcomplex II precedes assembly of the holocomplex, and thus increased proximity between EXOC5 and EXOC7 may reveal increased complex assembly required for proper exocyst function. Secondly, Ca^2+^-dependent remodeling of the plasma membrane may redirect plasma membrane-bound exocyst subunits, such as EXOC7, into closer proximity with other exocyst subunits. EXOC7 localization is dependent on synthesis and binding of PI(4,5)P_2_ (phosphatidylinositol 4,5-bisphosphate) (85). PI(4,5)P_2_ can be enriched within microdomains of the plasma membrane through a cascade starting with Ca^2+^-sensing by PKC, followed by activation of phosphatidylinositol-4-phosphate 5-kinase (PIP5K) (86, 87). From phosphatidylinositol 4-phosphate (PI4P), PIP5K synthesizes PI(4,5)P_2_, which it can be bound by PI(4,5)P_2_-interacting domains of EXOC7 (88). Through regulation of subunit assembly via direct phosphorylation and/or by reshaping plasma membrane domains, an increase in exocyst subunit proximity may signal the exocyst-mediated tethering of MVBs to secretion machinery on the plasma membrane and facilitating subsequent EV exocytosis observed during increased Ca^2+^ signaling.

Fourth, we found that associations between Rab27a and CD63^+^ MVBs are dynamically regulated during Ca^2+^-stimulated EV secretion. We used PLA to screen for MVB proximity with several Rab GTPases, such as Rab8a, Rab11a, Rab27a, Rab27b and Rab35, that hold roles in EV-related trafficking ranging from endosomal recycling to MVB trafficking (34, 89–92). We found that Rab27a exhibited the highest dynamic range with the PLA signal with high density in vehicle-treated cells, but dropping significantly after ionomycin treatment. This data is compatible with the well-documented and widely conserved role of Rab27a in the exocytic trafficking of exosome precursor-containing MVBs. Rab27a has been extensively implicated in the trafficking of EV precursor-containing MVBs across diverse cell types, with Rab27a downregulation and inhibition followed by deleterious consequences for small EV secretion (35, 36, 93, 94). Our findings suggest that Ca^2+^-stimulated EV secretion is accompanied by a reorganization of Rab27a associations with CD63^+^ MVBs. The reduction in Rab27a-CD63 proximity observed following ionomycin treatment may reflect dynamic remodeling of Rab27a interactions with CD63^+^ MVBs during stimulated secretion. As EV release is increased under these conditions, loss of PLA signal is consistent with progression of MVBs through membrane docking and fusion events that alter the spatial relationship between Rab27a and CD63-containing compartments (**Fig. 4A,C**).

Fifth, we utilized multiple orthogonal approaches to characterize the interaction between Rab27a and the exocyst complex. PLA analysis demonstrated that Ca^2+^ stimulation increased proximity between Rab27a and EXOC5 while coupling exocyst inhibition via ES2 with ionomycin treatment promoted an accumulation of Rab27a-associated CD63^+^ vesicles (consistent with impaired completion of MVB membrane delivery) (**Fig. 5A-C**). Through CoIP, we provide evidence of an interaction between the exocyst and Rab27a (**Fig. 6A**). While we were able to capture Rab27a with an EXOC4 immunocapture, the CoIP strategy faces challenges when discriminating between direct and indirect binding between Rab-effector complexes. Namely, the short-lived duration of these interactions is based on the rapid switching of Rab’s GTP-bound phase (51, 95). To experimentally determine the potential of direct protein-protein binding between Rab27a and EXOC6, we performed a solid-phase protein binding assay (**Fig. 6B**). Our study produced a concentration-dependent complex formation between purified Rab27a and EXOC6. This experimental evidence supports a claim for a direct interaction between the two proteins without additional quaternary structure provided by other exocyst subunits comprising the holocomplex. Interestingly, GTPγS- and GDPβS-loaded Rab27a produced distinct binding profiles, raising the possibility that EXOC6 association may be influenced by the nucleotide state of Rab27a. Although the nucleotide state of Rab27a could tune affinity with the exocyst, other biological aspects, including Ca^2+^ signaling, may also determine association during stimulated trafficking. We followed this experiment with molecular modeling to estimate energetic expressions relevant to protein-protein complex formation between Rab GTPases and the exocyst holocomplex (**Fig. 7A-D**). Although this modeling was performed in a computational setting without biochemically relevant post-translational modifications and nucleotide ligands, the similar dissociation constants and conserved binding topology between Rab GTPases and the exocyst suggest a potential direct interaction between Rab27a and exocyst complex subunits.

Proximity labeling-based proteomic analyses revealed that Ca²⁺ stimulation altered the composition of exocyst proteins recovered in Rab27a proximity-labeled proteomes. Notably the differential labeling of EXOC6 and EXOC6B suggests elevated intracellular Ca^2+^ promotes isoform-specific remodeling of exocyst subunits rather than simple recruitment of the Rab27a to the exocyst interface (**Fig. 8B**). Additional factors, including GTP deficiency, also influences exocyst proximity to Rab27a (**Fig. 8C**). Importantly, interpretation of the Rab27a^T23N^ proximity proteome is limited by the shorter biological half-life of this unstable dominant-negative mutant and its preferential GDP-bound state, which likely narrows its effector network (96). Thus, differences between the Rab27a^WT^ and Rab27a^T23N^ proximity proteomes likely reflect both altered nucleotide-dependent interactions and reduced mutant stability. Taken together, our experimental evidence supports the interpretation that Rab27a directly interacts with the exocyst in a dynamic manner during Ca^2+^ signaling within SkM myotubes mediating MVB trafficking and subsequent EV secretion.

Lastly, we found that the inhibition of AMPK, AKT, ERK1/2, CaMKII, and PKC results in differential changes of EV secretion revealed through our pHluorin-CD63-mScarlet expressing cell line (**Fig. 9E**). These kinases hold critical roles in reorganizing membrane structures, mobilizing contraction machinery and EV biogenesis in response to intracellular Ca^2+^ flux (20, 85, 97, 98). We observed that inhibition of AMPK, ERK1/2 and PKC in parallel with ionomycin treatment, significantly lowered ΔpHluorin-CD63 relative to ionomycin treatment alone. We posit that AMPK activation, which has been associated with aspects of vesicular and membrane dynamics, may play a distinct role in the upstream mobilization of EV biogenesis machinery via both Ca^2+^ and AMP sensing events. AMPK activation via CaMKII/Ca^2+^ sensing has been associated with increased actin remodeling alongside other aspects of vesicular and membrane dynamics (99, 100). During metabolic stress, however, AMPK can become activated in the presence of AMP and regulate lysosome homeostasis/MVB degradation via mTOR signaling (101). In this way, the biological context of AMPK activation may lead to differing outcomes relevant to EV release. Regarding the kinase AKT, pharmacological inhibition of AKT did not challenge ΔpHluorin-CD63 fluorescence versus ionomycin treatment alone. High levels of intracellular Ca^2+^ may overwhelm any contribution of AKT dynamics to EV secretion. We previously discussed the importance of Ca^2+^ signaling to activation of other kinases including ERK1/2, PKC and phosphoinositide-related kinases that promote exocyst function. We specifically highlight here the direct phosphorylation of exocyst subunits via both ERK1/2 and PKC. Exocyst activity can be positively or negatively regulated by kinases in order to meet the dynamic requirements of the cell. For example, during the arrest of cell growth between mitotic metaphase and anaphase, Cyclin Dependent kinases (CDK1) phosphorylate exocyst subunits to disrupt exocyst complex assembly and activity (19, 102). Contrasting the inhibition of the exocyst via CDK1, PKC and ERK/2 can both positively regulate exocyst activity through phosphorylation of EXOC2 and EXOC7(20, 21). We suggest the possibility of Ca^2+^ stimulation driving ERK1/2 and PKC activation (through either direct Ca^2+^ sensing or modulation by CaMKKβ) followed by downstream phosphorylation of subunits and the promotion of exocyst activity required during exosome secretion. Exploration of direct signaling-dependent regulation of the exocyst complex may more fully explain the interface between kinase activity and the modulation of trafficking and secretion networks.

The rapid expansion of EV research in the past two decades has allowed for the discovery of many exciting translational applications. New potential diagnostic and therapeutic uses of EVs have naturally captivated many scientists. Although this research is highly important, the field of EV research must not neglect the fundamental biology that drives the biogenesis and release of newly appreciated classes of EVs. Our understanding of the relationship between cell activity and EV release is defined by the activity of specific proteins belonging to signaling, trafficking and secretion families. Understanding the roles of these proteins allows us to define critical aspects of EV biology. The more clearly we describe the regulatory pieces of the biological puzzle, the more efficiently we can drive the successful generation of EV tools. Although the application of SkM-derived EVs in diagnostic and therapeutic settings remains appealing, we need to complement this work with a detailed characterization of the molecular biology underscoring the origin of bioactive EVs. Collectively, our data point to a Ca^2+^-dependent, modular association between Rab27a and the exocyst complex that organizes MVB trafficking and subsequent release of extracellular vesicles in cultured skeletal muscle cells. We posit that Ca^2+^ signaling is responsible for reshaping this molecular platform by tuning exocyst assembly and its recruitment to Rab27a. Lastly, we show that exocyst activity is necessary for EV secretion in response to ionomycin treatment, and this activity is likely regulated by Ca^2+^-sensing kinases PKC and ERK1/2, which are directly linked to the regulation of exocyst activity. Altogether, we describe signaling-dependent coordination among different arms of the secretion network that cooperate to promote the release of EVs in SkM cells in response to ionomycin-treatment-induced changes in intracellular calcium levels.

## EXPERIMENTAL PROCEDURES

### Cell Culture

WT C2C12 myoblasts (Sigma-Aldrich, 91031101-1VL) were grown in DMEM (Fisher Scientific 10-013-CV) supplemented with 10% FBS (Corning, MT35011CV), 1x Glutamax (ThermoScientific, 35050061), and 1x antimycotic/antibiotic (Corning, MT30004CI). Upon 100% confluence, myoblasts were differentiated using DMEM supplemented with 2% horse serum (Gibco, 26050-088), 1x Glutamax (ThermoScientific, 35050061), and 1x antimycotic/antibiotic (Fisher Scientific, MT30004CI) for 5 days. Cells were grown in an incubator (Isotemp, Fisher Scientific, 3532) with 5% CO_2_ at 37°C. Cells were passaged using Trypsin-EDTA (0.25%, sterile-filtered, BioReagent, suitable for cell culture, 2.5 g porcine trypsin and 0.2g EDTA, 4Na per liter of Hanks′ Balanced Salt Solution with phenol red) from Sigma-Aldrich (T4049). Preceding EV collection experiments, myotubes were incubated with DMEM containing 2% horse serum (> 24hr) that had been previously ultracentrifuged at > 100k xg; > 6hr to remove any serum-derived EVs. Multinucleated C2C12 myotube morphology was validated using DAPI imaging following differentiation. Cells were also tested for *Mycoplasma* using PCR provided with primers a positive control template (Abcam, ab289834).

### Co-Immuno Precipitation

Differentiated WT C2C12 myotubes were harvested and lysed using CoIP buffer (20 mM Tris-HCl, pH 8.0, 150 mM NaCl, 2mM EDTA, and 1% Triton X-100) on ice for 30 minutes. Cell debris was pelleted at 16k x g for 15 minutes at 4°C and supernatant was pre-cleared using Pierce Protein A/G Magnetic Beads (ThermoFischer, 88802). After lysate was incubated for 1 hour at 4°C on rotation, pre-cleared lysate was divided into 3 equal volume fractions. The first fraction was incubated with no IgG. The second fraction was incubated with isotype IgG (Mouse) at ∼1μg IgG/mg lysate. The third fraction was incubated with anti-EXOC4 IgG. All three fractions were rotated at 4°C for 16 hours. After antibody incubation, magnetic beads were introduced to the lysate fractions and rotated at 25°C for 1 hour. Supernatants were removed from magnetic beads and beads were washed three times with CoIP buffer before being boiled in loading buffer for protein gel electrophoresis.

### Plasmid cloning and sequencing

Plasmids were constructed using primers ordered via Integrated DNA Technologies (IDT). Gibson assembly kits (New England Biolabs, E5510S) were used to assemble DNA fragments into novel plasmids. Trangene-containing plasmids were transformed using chemically competent NEB 5-alpha *E. coli* cells and were grown on Kanamycin (50μg/mL) or Carbenicillin (100μg/mL) agar plates. Whole plasmid DNA was amplified before being sequenced via the Oxford Nanopore technique through Plasmidsaurus.

### Transfections

C2C12 myoblasts were transfected using the Lipofectamine 3k kit (Thermo Scientific, L3000008) at approximately 70% confluence. Upon incubation of cells with plasmid-reagent mix for 24 hours, transfection-media mixture was removed and media containing a final concentration of 0.2mg/mL Hygromycin B (Sigma-Aldrich, H7772-100MG) was supplemented to select for transgene-expressing C2C12 cells.

### PLA

For Proximity Ligation Assay (PLA), C2C12 myotubes were grown on #1.5 glass cover slips. Cells were treated and washed with cold PBS twice before fixation (4% PFA, 15 minutes at room temperature). PFA was removed from fixed cells and again washed with 1x PBS twice. Cells were then permeabilized with 0.1% Triton-X 100 in PBS for 15 minutes on rotation at room temperature. Permeabilization buffer was removed washed three times with 1x PBS. Duolink blocking solution (∼40 μL/cover slip) was then added to fixed cells overnight at 4°C on rotation. Primary antibodies against protein targets were validated in advance through immunocytochemistry and confocal imaging in advance of PLA usage. Antibodies were then diluted in Duolink antibody diluent solution 1:100 to a final volume of ∼40 μL/cover slip and incubated with parafilm covering glass cover slip overnight at 4°C. The antibody dilution was then aspirated, and coverslip was washed with 1x Wash Buffer A 2x 5 minutes. A 40 μL dilution of PLA probe MINUS and PLA probe PLUS was incubated on sample for 1 hour at 37°C in a humidity chamber. Probes were ligated using a dilution of Duolink ligase in ligation buffer and incubating on samples for 30 minutes at 37°C in a humidity chamber. Samples were again washed with Wash Buffer A then incubated with Duolink polymerase in Amplification buffer for 100 minutes at 37°C in a humidity chamber. Polymerase reaction buffer was aspirated, and samples were washed twice with 1x Duolink Wash Buffer B before a final wash in 0.01x Wash Buffer B for 1 minute. Slides were mounted using Duolink mounting media with DAPI and sealed using nail polish. PLA puncta were quantified using a macro in ImageJ and normalized to the number of nuclei within each field to account for differences in cell density between imaging fields.

### Fluorescent microscopy imaging

For PLA quanitification and subsequent analysis, twenty-one fields of view were counted for each experimental condition. Fluorescent images (PLA and immunofluorescent images) taken for quantification were collected on the epifluorescent microscope (Olympus BX41). Representative images were additionally captured using a Leica SP8 and Leica Thunder Live Cell 3D microscope.

### FACS

Single C2C12 cells expressing the pHluorin-CD63-mScarlet transgene isolated using the BD FACS Aria IIu. to control for viability, Draq7 was incubated with resuspended C2C12 myoblasts; cells positive for mScarlet alone were used as a fluorescent positive control for gating.

### Small Molecule Inhibitors

Compounds including Dantrolene [Cayman Chemical Company, 14326] (10 μM, 12 hr), Manumycin A [Cayman Chemical Company, 10010497] (10 μM, 12 hr), Endosidin 2 [Cayman Chemical Company, 21888] (30 μM, 30 min), AKT Inh. VIII (1 μM, 12 hr) [Chemical Company, 14870], BAY3827 [TargetMol, T73350] (1 μM, 12 hr), KN-93 [Selleck Chemicals, S6787] (1 μM, 12 hr) and SCH772984 [TargetMol, T6066] (5 μM, 12 hr) were incubated with pHluorin-CD63-mScarlet C2C12 myotubes to perform inhibition and challenged with Ionomycin.

### Fluorescence platereader assays

Changes in pHluorin fluorescence were measured using the Varioskan LUX Multimode Microplate Reader. Fluorescent kinetics were monitored with ex/em 488 nm/511 nm and recorded every 5 minutes at 37°C.

### Calcium influx assay

To measure relative changes of intra- and extracellular Ca^2+^ in C2C12 myotubes, cells were seeded and differentiated in 96 well plates (Cellvis, P96-1.5P). Cells were washed with 1x PBS before being incubated with 5 μM FURA-2AM (Cayman Chemical Company, 14591) in Fluorobrite media (Thermo Scientific, A1896701) for 1 hour at 37°C. Cells were then washed and treated with 0-20 μM ionomycin (Cayman Chemical Company, 10004974) in Fluorobrite media. Fluorescence measurements at excitation wavelengths 340 nm and 380 nm and emission at 510 nm were taken every 5 minutes. Data was fit to a non-linear exponential model (Y=Bmax*X/(Kd+X) + NS*X + Background).

### Cell Viability Assay

C2C12 myotubes were seeded at a density of ∼1.0x10^6^ cells/96-well plate and differentiated once cells reached 100% confluence. Cells were washed with 1x PBS to remove serum. Media was replaced with Fluorobrite media before incubating with LIVE/DEAD ® Viability/Cytotoxicity Kit *for mammalian cells* (FisherScientific L3224). Fluorescence was monitored at ex/em 495 nm/515 nm (Calcein) and ex/em 495 nm/635 nm (EthD-1) over 30 minutes at 37°C.

### EV isolation and characterization

C2C12 myotubes were grown and differentiated in 6-well plates with glass #1.5 coverslips. After small molecule treatment of C2C12 myotubes, conditioned cell media was collected and spun at 3k x g for 20 minutes at 4°C to pellet cell debris (Eppendorf 5804R). C2C12 myotubes were then fixed for 20 minutes at room temperature with 4% paraformaldehyde before washing with PBS for downstream nuclei quantification. The conditioned cell media supernatant was then transferred to Amicon Ultra-4 centrifugal filters (UFC801096) and spun at 3k x g for 20 minutes at 4°C to concentrate conditioned cell media. Upon concentrating volumes to 150 μL, concentrates were spun at 16k x g at 4°C for 15 minutes to clear any aggregates or debris. Cleared supernatant volumes were adjusted to 150 μL using 1x PBS and loaded onto qEV 35 nm single size exclusion chromatography (SEC) columns (IZON: ICS-35) and fractionated using automated fraction collectors (AFC) (IZON, AFC-V2). Mobile phase for SEC was 0.22 μm filtered 2x PBS (274 mM; pH 7.4). After a void volume of 0.87mL had passed through the AFC, a single fraction with a volume of 0.97mL was collected per replicate of concentrated conditioned cell media. Isolated particles were stored temporarily at 4°C for up to 48hr before size and concentration analysis were performed.

### TRPS measurements

Particle suspension was diluted 1:3 in 2x PBS (0.22 μm filtered; 274 mM; pH 7.4). In advance of sample measurements, the Exoid V1 (IZON) was fitted with a nanopore NP100 and characterized after nanopore coating and wetting procedures as recommended by the manufacturer. The nanopore NP100 was then calibrated with CPC100 (100 nm) beads at a dilution of 1:2000. To measure sample size and concentrations, 35μL of each sample was reverse pipetted onto nanopore NP-100 and recorded at 3 calibration pressures (400, 600, 800 Pa) with a voltage at ∼140 nA and RMS of < 10 pA. New nanopores were coated, calibrated and exchanged after every 4^th^ sample measurement to maintain high measurement fidelity.

### EV protein concentration

To perform a Western blot analysis for EV markers and cargo, 200 μL of particle suspension was centrifuged in Amicon ultra centrifugal 3kDa MWCO filters (Millipore Sigma, UFC5003) at 16k x g at 4°C for 30 minutes. When the sample volumes approached 20 μL (concentration at 10x), the samples were resuspended in 2x SDS loading buffer (Biorad, 161-0737) and denatured at 95°C for 5 minutes. Parental cell lysate (C2C12 myotube) was run in parallel with isolated EV material as a control.

### Western blotting

Resuspended EV protein samples were loaded onto Mini-PROTEAN TGX precast gels (4-15% acrylamide) (Bio-Rad, 4561085) and run at 150V for 45 minutes. Western blotting was performed using transfer of protein form SDS page gel onto Immun-Blot LF PVDF membrane (BioRad, 1620264) using Trans-Blot Turbo system (BioRad, 1704150) for 2.5A, 25V, 13 minutes. Total protein transferred into the membrane was resolved using Ponceau S solution (sigmaaldrich, P7170-1L) or Revert 700 nm Total Protein Stain (LiCor, 926-11011) before being blocked for 1 hour at room temperature (LiCor Intercept, 927-70001). The membrane was then probed with primary antibodies at a dilution of 1:1000 overnight at 4°C. Primary antibodies were removed and membranes were washed for 5 minutes with 1x PBS 3 times before being incubated with fluorescent secondary antibodies (IRDye 680RD Donkey anti-Mouse, LiCor, 926-68072; IRDye 800CW Goat anti-Rabbit, LiCor, 926-32211) at 1:10000 dilution for 45 minutes at room temperature. Membranes were washed 3x for 5 minutes each with PBST before being incubated in PBS for imaging. Membranes were imaged using the LiCor Odyssey F IR scanner. Images were opened and saved as TIFFs via ImageStudio software. For each band, a rectangular region of interest (ROI) was drawn around the band, and the median pixel intensity was recorded to quantify bband intensity. Background signal was measured using an equivalent rectangular ROI in an area adjacent to the band and subtracted from the corresponding band measurement to obtain the background-corrected signal.

### Nano Flow Cytometry

Post-SEC isolated EVs from WT and pHluorin-CD63-mScarlet C2C12 myotubes were measured on a NanoFCM Nanoanalyzer featuring fluorescence detection of side scatter (523/14 nm) representing total particles and PE (580/40 nm) representing mScarlet^+^ EVs.

### TEM

Galvanized copper Formvar-coated grids (Electron Microscopy Sciences, FCF200-CU-50 type SB) were applied with 4μL with EV sample for 45 seconds before being wicked off the grid. Distilled water (10 μL) was used to wash sample before being wicked again. Sample was stained (2% Uranyl Acetate; 4μL) for 45 seconds before being wicked. Sample was again washed with 10 μL distilled water. Samples were imaged using a Hitachi HT7700 Transmission Electron Microscope (120kV) paired with an AMT BioSprint16M-ActiveVu 16 megapixel cooled 4896 x 3920 pixel CCD camera.

### Modeling protein-protein interactions

The structure for full length, monomeric Rab8a, Rab10, Rab11a and Rab27a were modeled using AlphaFold 3’s server platform (41). Each structure was first loaded into Pymol™ 2.4.2 where the structure was protonated. The structure for the exocyst complex (PDB 5YFP (18)) was also loaded into Pymol and subsequently protonated after removal of solvent and organic groups. As an octamer with 8 individual subunits, a continuous single chain was iterated to represent all subunits as a single amino acid range. To model the exocyst-Rab GTPase complexes, the Rab GTPase, IgG Fc Domain (PDB 4QGT (40)) and exocyst PDBs were then saved and exported to the HADDOCK 2.4 protein modeling platform (103, 104). Buried residues were included in modeling selection. Non-polar hydrogens were removed, DNA/RNA restraints were used and the center of mass was defined using restraints to enforce contact between the molecules. Random exclusion was used to exclude a fraction of the ambiguous restraints (AIRs). After modeling was performed via HADDOCK 2.4, X,Y,Z coordinates were extracted from the PDB models and inserted into a dataframe via R (4.4.1). The free energy of complex formation on the all models were assessed using HADDOCK PRODIGY web-based platform (105). All energetic simulations were performed at 37°C for each of the selected protein complex models and produced K_D_ equilibrium constants from 4.6e-6 – 2.7e-7 M for exocyst-Rab27a complexes.

### Solid-phase Protein Binding Assay

96-well plates coated with mouse-produced anti-Flag Tag antibodies (GenScript, L00455C) were coated with 150ng/well human recombinant EXOC6-Flag Tag (BosterBio, PROTQ8TAG9) overnight at 4°C. The 96-well plate was washed with 0.22μm filtered PBS before blocking for 1hr at 25°C (LiCor, 927-70001). Purified 6xHis-Rab27a (LSBio, LS-G779) was first chelated with 1mM EDTA for 20 minutes at 25°C to allow for exchange of either 10 mM GTPγS (Sigma, 20-176, G7637) or 10 mM GDPβS (Sigma, G7637). 100 mM MgCl_2_ was then added to stabilize the nucleotide binding with Rab27a. Rab27a, purified BSA (Thermo, 3235), and 6xHis-GFP (SinoBiological, 69001-S08E) were then titrated at 5000, 2500, 1000, 750, 500, 100, 20, 0 nM. Concentration of purified EXOC5-GST (LSBio, LS-G32272) with use of microcentrifuge filters (3,000 NMWL (nominal molecular weight limit), Microcon, 42404) was required to reach 1000, 500, 200, 50, 15, 5 nM concentrations. The concentration of EXOC5-GST was verified via a Nanodrop 280 nm (ThermoFisher, ND-ONE-W) = given the extinction coefficient (42385M^-1^ cm^-1^). Rab27a, BSA, GFP, and EXOC5 were incubated overnight at 4°C before washing (PBS pH 7.6, 2 mM MgCl_2_, 0.05% Tween-20) and then incubated with a 1:500 dilution of anti-Rab27a primary antibody overnight at 4°C (Proteintech, 17817-1-AP) in Rab27a wells and BSA wells, and anti-EXOC5 (Invitrogen, PA5-109698) was incubated in EXOC5 wells. Intrinsic fluorescence of GFP (488 nm/520 nm) was used for GFP readout. All wells were then washed 3x before incubation with 1:5000 anti-Rabbit 488 nm (DyLight, I488) at 25°C for 1 hour. The plate was then washed 3x with 1x PBS before imaging on the Varioskan LUX Multimode Microplate Reader (Thermo, VLBLATD0). Analysis of data was performed using the following model in Prism Graphpad: Y=Bmax*X/(Kd+X) + NS*X + Background.

### Biotin labeling and streptavidin pulldown

Four independent biological experiments were performed on four separate days, with each experiment including all eight experimental conditions, yielding four biological replicates per condition (n = 4). Transgenic, UltraID-expressing C2C12 myotubes were differentiated in 150x30 mm dishes using DMEM (Thermo Scientific, 31053028) supplemented with 1x final concentration of Glutamax (Gibco, 35050061), 2% horse serum (Gibco, 26050088) and 0.2 mg/mL Hygromycin B (Sigma-Aldrich, 31282-04-9). 48 hr prior to treatment with biotin, myotubes were incubated with media supplemented with dialyzed horse serum to reach a final concentration of 2% (20mL horse serum dialyzed overnight in 2L of 1x PBS at 4°C within SpectraPor dialysis tubing (132720)) to quench any source of biotin before labeling. On treatment day, cells were washed with warm 1x PBS (∼15mL /dish) before being incubated with media [DMEM (Thermo Scientific, 31053028) supplemented with 1x final concentration of Glutamax (Gibco, 35050061), 2% dialyzed horse serum (Gibco, 26050088) and 0.2mg/mL Hygromycin B (Sigma-Aldrich, 31282-04-9) additionally supplemented with 50 μM biotin (Millipore Sigma, 58-85-5) and either vehicle (EtOH) or Ionomycin (10 μM) for 15 minutes at 37°C. Cells were then removed from the incubator, washed 2x with cold 1x PBS, and incubated with 400 μL RIPA (ThermoFisher, 89900) with 1x protease inhibitor (Millipore Sigma, P8340-1ML) and 1x phosphatase inhibitor (ThermoScientific, PI78420) and scraped into a 2mL Eppendorf tubes. Lysates were rotated at 4°C for 30 minutes before insoluble fractions were pelleted at 16k RCF at 4°C. Soluble fractions were normalized by volume to 2.5mL with 1x PBS before loading through PD-10 G25 desalting columns to remove excess biotin (Cytiva, 17-0851-01) and eluted with 3.5mL PBS. Protein yields were normalized by concentration and incubated with Neutravidin-agarose 50% slurry (ThermoScientific, 29201) for 2 hours at 25°C on rotation. Slurries were washed with RIPA Buffer 3x before additional washes in 1x PBS. Two final washes with NaHCO_3_, pH 7.5 buffer were performed before beads were pelleted and frozen at -80°C for storage.

### Proteomics

After NeutrAvidin pulldown of biotinylated proteins, bead slurries were sent to the University of Arkansas National Resource for Quantitative Proteomics for sample preparation and data acquisition. Each biological sample was analyzed by a single LC-MS injection. A pooled sample was injected at the beginning, middle, and end of the analytical run as a quality-control sample. Protein samples were reduced, alkylated, and digested using filter-aided sample preparation (FASP). On-bead protein samples were supplemented with 200 μL of 100 mM triethylammonium bicarbonate (TEAB) and reduced with 10 μL of 200 mM tris(2-carboxyethyl)phosphine (TCEP) for 30 min at 37°C. Samples were subsequently transferred to FASP filter units, centrifuged at 14,000 × g for 20 min, and alkylated with 100 μL of 30 mM iodoacetamide (IAA) in TEAB for 20 min in the dark. Tryptic digestion was performed using sequencing-grade modified porcine trypsin (Promega) at a 1:50 trypsin-to-protein ratio. Samples were incubated in a humidified chamber at 47°C for 18 h for a single overnight digestion (106). Following digestion, samples were centrifuged at 14,000 × g for 10 min, and 80 μL of 100 mM TEAB was added to the filter units before a second centrifugation at 14,000 × g for 10 min. Peptide-containing filtrates were acidified to 0.5% formic acid/0.1% trifluoroacetic acid using 5% formic acid/1% trifluoroacetic acid and desalted using a Sep-Pak 96-well plate. Desalted peptides were subsequently filtered and dried by SpeedVac prior to LC-MS/MS analysis. No offline peptide fractionation was performed. Tryptic peptides were separated by reverse-phase chromatography using XSelect CSH C18 2.5 μm resin (Waters) on an in-line 150 × 0.075 mm column using an UltiMate 3000 RSLCnano system (Thermo Scientific) at a flow rate of 300 nL/min and an ambient column temperature of 22°C. Peptides were eluted using a 60-min gradient from a 98:2 to 65:35 buffer A:buffer B ratio. Buffer A consisted of 0.1% formic acid and 0.5% acetonitrile in water, and buffer B consisted of 0.1% formic acid and 99.9% acetonitrile. Eluted peptides were ionized by electrospray at 2.4 kV through a heated capillary maintained at 275°C and analyzed using an Orbitrap Exploris 480 mass spectrometer (Thermo Scientific). Precursor spectra were acquired from 385–1015 Th at a resolution of 60,000, with 100% AGC, a maximum injection time of 50 ms, and an RF parameter of 40%. DIA was performed in positive ion mode using an approximately 3-s cycle time. The DIA method used 40 × 15 Th isolation windows spanning 394–1009 Th, with a resolution of 15,000, a normalized AGC target of 500%, a maximum injection time of 40 ms, and HCD collision energy set to a normalized value of 30%. A staggered 15 Th window pattern with optimized window placements spanning 400–1015 Th was incorporated into the DIA acquisition.

Following data acquisition, raw data were analyzed using Spectronaut (Biognosys, version 20.3) using the DirectDIA workflow against the UniProt *Mus musculus* reference proteome (Proteome ID: UP000000589; Taxon ID: 10090; fourth version of 2025). Label-free protein quantification was performed using MS2-level quantities. Protein identification and quantification used a precursor and protein q-value cutoff of 1%, with decoy generation enabled. The protein inference workflow was set to Quant 2.0 with the IDPicker inference algorithm, and protein-group quantities were calculated as the median of peptide and precursor quantities. Up to two missed cleavages were allowed. As a provisional value pending confirmation from the proteomics core, precursor and fragment mass tolerances were set to 9.2 ppm and 22.4 ppm, respectively. A minimum of one unique peptide was required for protein-group identification, and unique and razor peptides were used for quantification. Carbamidomethylation of cysteine was specified as a fixed modification, while protein N-terminal acetylation and methionine oxidation were specified as variable modifications. Protein MS2 intensity values were evaluated for quality using ProteiNorm (107), incorporated within proteoDA version 1.0. Raw protein intensity data were evaluated using multiple normalization strategies, including log2 transformation alone, median normalization, mean normalization, variance-stabilizing normalization (VSN), quantile normalization, cyclic loess normalization, robust linear regression normalization (RLR), and global intensity normalization (GI). Performance was assessed using intragroup correlation, intensity distribution consistency, pooled coefficient of variation (PCV), pooled median absolute deviation (PMAD), pooled estimate of variance (PEV), principal component analysis (PCA), and hierarchical clustering (**Supp. S4**). Statistical analyses were performed using proteoDA version 1.0 and the limma framework with empirical Bayes (eBayes) smoothing of standard errors (108). Data were log2 transformed prior to statistical analysis. Proteins with an FDR-adjusted p-value < 0.05 and a fold change > 2 were considered significant. Data visualization and processing were performed using R version 4.4.1 (R Core Team, 2024), with the tidyverse, ggplot2, and scales packages. The tidyverse was used for data import and manipulation, ggplot2 for data visualization, and scales for continuous scale formatting (109–111). Proteomics results are available within ProteomeXchange Consortium via the PRIDE (public repository of protein and peptide identifications) partner repository with the dataset identifier PXD077110.

### Quantification

For quantified data, statistical analyses were performed using the replicate structure indicated in the corresponding figure legends. Experiments were independently repeated at least three times where indicated, while selected representative experiments containing technical replicate wells were analyzed separately as indicated in the figure legends. For pairwise comparisons, data were assessed for normality using the Shapiro–Wilk test, followed by unpaired t tests for normally distributed data or Mann–Whitney U tests for non-normally distributed data. For experiments involving multiple treatment groups, one-way or two-way ANOVA or Kruskal–Wallis tests were used as appropriate. ANOVA analyses were followed by Tukey’s multiple-comparisons tests or uncorrected Fisher’s LSD tests, as indicated in the corresponding figure legends. Kruskal–Wallis analyses were followed by Dunn’s multiple-comparisons tests. For two-way ANOVA, main effects and interaction effects were evaluated, followed by post hoc comparisons where appropriate. Potential outliers were identified using an interquartile range (IQR)-based approach. The first (Q1) and third (Q3) quartiles were calculated, and the IQR was defined as Q3 − Q1. Lower and upper limits were calculated as Q1 − 1.5 × IQR and Q3 + 1.5 × IQR, respectively. Observations falling below the lower limit or above the upper limit were considered potential outliers.

### Antibodies

Antibodies for PLA were previously validated with immune-cytochemistry and western blotting. Antibodies used in western blotting were validated using lysate controls that matched manufacturers published banding patterns.

AKT (Pan) [Cell Signaling, 40D4]

AMPK alpha-1 [Invitrogen, A5-15815]

AS160 [Invitrogen, MA5-14840]

Calnexin [Proteintech, 66903-1-IG]

CD9 [Abcam, ab275018]

CD63 [Proteintech, 67605-1-Ig]

CD81 [Proteintech, 66866-1-Ig]

GFP [Santa Cruz, sc-9996]

Dystrophin [Santa Cruz, sc-73592]

ERK1/2 (p44/42) [Cell Signaling, 4695]

EXOC5 [Santa Cruz, sc-514802]

EXOC5 [Invitrogen, PA5-109698]

EXOC7 [Santa Cruz, sc-365825]

GAPDH [Proteintech, 60004-1-Ig]

Histone H3 [Proteintech, 17168-1-AP]

IgG Mouse Normal [Santa Cruz, sc-2025]

IRDye 680RD Donkey anti-Mouse [LiCor, 926-68072]

IRDye 800CW Goat anti-Rabbit [LiCor, 926-32211]

Myc Tag [Proteintech, 66004-1-AP]

Phospho-AKT (pSer473) [Cell Signaling, 9271S]

Phospho-AMPK (pThr172) [Cell Signaling, 50081]

Phospho-AS160 (pThr642) [Invitrogen, 44-1071G]

Phospho-ERK1/2 (pT202/pY204) [Santa Cruz, sc-136521]

RAB8 [Proteintech, 55296-1-AP]

RAB11 [Cell Signaling, 5589S]

RAB27A [Proteintech, 17817-1-AP]

RAB27B [Proteintech, 13412-1-AP]

RAB35 [Proteintech, 11329-2-AP]

rSEC8 (EXOC4) [ENZO, ADI-VAM-SV016]

Streptavidin (800 nm) [LiCor, 926-32230]

SYTL4 [Proteintech, 12128-1-AP]

TSG101 [Proteintech, 28283-1-AP]

## DATA AVAILABILITY

The mass spectrometry proteomics data have been deposited to the ProteomeXchange Consortium via the PRIDE partner repository with the dataset identifier PXD077110. Western blot images are shown in supplemental materials. All PLA, molecular modeling, and fluorescent plate reader data are available upon request. Plasmids and corresponding whole plasmid sequencing maps were deposited to Addgene and are available for purchase.

## SUPPORTING INFORMATION

This article contains supporting information.

## AUTHOR CONTRIBUTIONS

This study was conceived by Luke T. Nelson and Noemi Polgar. L.T.N. executed experiments, including data collection and analysis. L.T.N., Jeremy Boulestreau, and N.P. interpreted the results and provided key insights. All authors were involved in editing the manuscript and approving the final submitted version, which was written by L.T.N., and N.P. All authors have read and agreed to the published version of the manuscript.

## FUNDING AND ADDITIONAL INFORMATION

We would like to acknowledge several core facilities and their funding support that allowed for our use of specialized equipment including the use of the University of Hawaii Molecular and Cellular Immunology Core’s BD FACS Aria Iiu for FACS (P30GM114737 from the Centers of Biomedical Research Excellence (COBRE) program of the NIGMS (National Institute of General Medical Sciences), the University of Hawaii Cancer Center’s Microscopy, Imaging, and Flow Cytometry Equipment Core Facility’s Leica Thunder Live Cell 3D microscope for fluorescent imaging (NIH 1S10OD028515-01), and the University of Hawaii and Pacific Biosciences Research Center’s Biological Electron Microscopy Facility for their training and use of the Hitachi HT7700 Transmission Electron Microscope for TEM imaging (NSF DBI-1040548). The support from the NIMHD (National Institute on Minority Health Disparities) has also allowed us to acquire the NanoFCM (N30E) for nano-flow cytometry applications (NIMHD Ola Hawaii U54MD007601). We would also like to acknowledge the IdeA (Institutional Development Award) National Resource for Quantitative Proteomics at University of Arkansas and the support from NIGMS (National Institute of General Medical Sciences) (R24GM137786) for proteomics and bioinformatics services. The project was also supported by funds for the Diabetes Research Center (NIGMS P20GM113134). The content is solely the responsibility of the authors and does not necessarily represent the official views of the National Institutes of Health.

## Abbreviations

AKT: Protein Kinase B
AMPK: AMP-activated Protein Kinase
AS160: Akt Substrate of 160 kDa
C2C12: subclone 12 derived from the original mouse myoblast
C2 cell line Ca^2+^: Calcium
CaMKII: Calcium/Calmodulin-dependent Protein Kinase II
CD63: Cluster of Differentiation 63
CoIP: Co-Immuno Precipitation
ER: Endoplasmic Reticulum
ERK1/2: Extracellular signal-Regulated Kinases 1 and 2
ES2: Endosidin 2
EV: Extracellular Vesicle
EXOC: Exocyst
FURA 2AM: Fura-2-acetoxymethyl ester
GAP: GTPase Activating Protein
GDI: Guanosine Dissociation Inhibitor
GDP: Guanosine Diphosphate
GEF: Guanosine Exchange Factor
GGTase: Geranyl-Gernanyl Transferase
GTP: Guanosine Triphosphate
Iono: Ionomycin
KD: Dissociation Constant / Equilibrium Constant
LCMS: Liquid Chromatography Mass Spectrometry
MVB: Multivesicular Body
PI: Phosphoinositide
Piggybac: Transposon identified in *Trichoplusia ni* that can insert into baculovirus
PLA: Proximity Ligation Assay
Rab GTPase: Ras-like protein from brain GTPase
SEC: Size Exclusion Chromatography
SkM: Skeletal Muscle
SNARE: Soluble N-ethylmaleimide-Sensitive Factor Attachment Protein Receptors
SR: Sarcoplasmic Reticulum
SYTL: Synaptotagmin-like Protein
UltraID: Evolved hyperactive mutant of BioID Biotin Ligase

## SUPPLEMENTAL MATERIAL

### Primers

NameSequence (5’-3’)

BioID L39P F1:GGCAGAGGCGGCCCGGGCAGAAAGTGGCTGAGC

BioID L39P R1: CTTGGTCTGTCTGTCGGCCAC

BioID L39P F2: GAGGCGGCCCGGGCAGAAAGTGGCTGAGCCA

BioID L39P R2: TGCCCTTGGTCTGTCTGTCGG

Start Methionine Q5 F: GCCACCATGGCGGCAGAAGCAG

Start Methionine Q5 R: GGTACCTGAATTCTTTGCCAAAATGATGC

BioID Insert F: CATCATTTTGGCAAAGAATTCAGGTACCGCCACCATGGAACAAAAACTCATCTCAGAAGAGGATCTCG

BioID Insert R: CAGACATAGAATTCGAAGCTTGAGCTGGCGGCCGCCCCTTCTCCTTGAACT

Rab27a Insert F: AGGGGCGGCCGCCAGCTCAAGCTTCGAATTCTATGTCTG

Rab27a Insert R: AACAACAATTGCCCTTCTAGGATCCCTCAACAGCCACATGCCCC

Exoc4 F: CATCATTTTGGCAAAGAATTCAGGTACCCCATGGCGGCAGAAGCAGCTG

Exoc4 R: AACGGTAGTTATCTTCTTGTCCTTGGTGGC

Exoc4-BioID F: GCCACCAAGGACAAGAAGATAACTACCGTTATGGAACAAAAACTCATCTCAGAAGAGGATC

Exoc4 BioID R: AACAACAATTGCCCTTCTAGGATCCTCATGGCGGCCGCCCCTTCTCCTTGAACT

BioID F: GACTGAAGGAGTGGAACG

BioID R: CGTTCCACTCCTTCAGTC Rab27a R: CCTGCTGTGTCCCATAAC

Exoc4 F: CACACGAACACGGATCTCC

Rab27a Q5 T23N Forward: GTGTAGGGAAGAACAGTGTACTTTACCAATATACAGATGG

Rab27a Q5 T23N Reverse: CAGAGTCTCCCAAAGCTAAAAA

## REFERENCES

1. Aminzadeh, M. A., Rogers, R. G., Fournier, M., Tobin, R. E., Guan, X., Childers, M. K., Andres, A. M., Taylor, D. J., Ibrahim, A., Ding, X., Torrente, A., Goldhaber, J. M., Lewis, M., Gottlieb, R. A., Victor, R. A., and Marbán, E. (2018) Exosome-Mediated Benefits of Cell Therapy in Mouse and Human Models of Duchenne Muscular Dystrophy. Stem Cell Rep. 10, 942–955

2. Castaño, C., Mirasierra, M., Vallejo, M., Novials, A., and Párrizas, M. (2020) Delivery of muscle-derived exosomal miRNAs induced by HIIT improves insulin sensitivity through down-regulation of hepatic FoxO1 in mice. Proc. Natl. Acad. Sci. 117, 30335–30343

3. Bei, Y., Xu, T., Lv, D., Yu, P., Xu, J., Che, L., Das, A., Tigges, J., Toxavidis, V., Ghiran, I., Shah, R., Li, Y., Zhang, Y., Das, S., and Xiao, J. (2017) Exercise-induced circulating extracellular vesicles protect against cardiac ischemia–reperfusion injury. Basic Res. Cardiol. 112, 38

4. Wang, Y., Liu, Y., Zhang, S., Li, N., Xing, C., Wang, C., Wang, J., Wei, M., Yang, G., and Yuan, L. (2023) Exercise Improves Metabolism and Alleviates Atherosclerosis via Muscle-Derived Extracellular Vesicles. Aging Dis. 14, 952

5. Kargl, C. K., Jia, Z., Shera, D. A., Sullivan, B. P., Burton, L. C., Kim, K. H., Nie, Y., Hubal, M. J., Shannahan, J. H., Kuang, S., and Gavin, T. P. (2023) Angiogenic potential of skeletal muscle derived extracellular vesicles differs between oxidative and glycolytic muscle tissue in mice. Sci. Rep. 13, 18943

6. Estrada, A. L., Valenti, Z. J., Hehn, G., Amorese, A. J., Williams, N. S., Balestrieri, N. P., Deighan, C., Allen, C. P., Spangenburg, E. E., Kruh-Garcia, N. A., and Lark, D. S. (2022) Extracellular vesicle secretion is tissue-dependent ex vivo and skeletal muscle myofiber extracellular vesicles reach the circulation in vivo. Am. J. Physiol.-Cell Physiol. 322, C246–C259

7. Van Pelt, D. W., Vechetti, I. J., Lawrence, M. M., Van Pelt, K. L., Patel, P., Miller, B. F., Butterfield, T. A., and Dupont-Versteegden, E. E. (2020) Serum extracellular vesicle miR-203a-3p content is associated with skeletal muscle mass and protein turnover during disuse atrophy and regrowth. Am. J. Physiol.-Cell Physiol. 319, C419–C431

8. Théry, C., Zitvogel, L., and Amigorena, S. (2002) Exosomes: composition, biogenesis and function. Nat. Rev. Immunol. 2, 569–579

9. Joshi, B. S., De Beer, M. A., Giepmans, B. N. G., and Zuhorn, I. S. (2020) Endocytosis of Extracellular Vesicles and Release of Their Cargo from Endosomes. ACS Nano. 14, 4444–4455

10. Gillingham, A. K., Sinka, R., Torres, I. L., Lilley, K. S., and Munro, S. (2014) Toward a Comprehensive Map of the Effectors of Rab GTPases. Dev. Cell. 31, 358–373

11. Grosshans, B. L., Ortiz, D., and Novick, P. (2006) Rabs and their effectors: Achieving specificity in membrane traffic. Proc. Natl. Acad. Sci. 103, 11821–11827

12. Knödler, A., Feng, S., Zhang, J., Zhang, X., Das, A., Peränen, J., and Guo, W. (2010) Coordination of Rab8 and Rab11 in primary ciliogenesis. Proc. Natl. Acad. Sci. 107, 6346–6351

13. Bhuin, T., and Roy, J. K. (2019) Developmental expression, co-localization and genetic interaction of exocyst component Sec15 with Rab11 during Drosophila development. Exp. Cell Res. 381, 94–104

14. Wu, H., Rossi, G., and Brennwald, P. (2008) The ghost in the machine: small GTPases as spatial regulators of exocytosis. Trends Cell Biol. 18, 397–404

15. Bai, S., Hou, W., Yao, Y., Meng, J., Wei, Y., Hu, F., Hu, X., Wu, J., Zhang, N., Xu, R., Tian, F., Wang, B., Liao, H., Du, Y., Fang, H., He, W., Liu, Y., Shen, B., and Du, J. (2022) Exocyst controls exosome biogenesis via Rab11a. Mol. Ther. Nucleic Acids. 27, 535–546

16. Liu, D.-A., Tao, K., Wu, B., Yu, Z., Szczepaniak, M., Rames, M., Yang, C., Svitkina, T., Zhu, Y., Xu, F., Nan, X., and Guo, W. (2023) A phosphoinositide switch mediates exocyst recruitment to multivesicular endosomes for exosome secretion. Nat. Commun. 14, 6883

17. Lee, C., Lepore, D., Lee, S.-H., Kim, T. G., Buwa, N., Lee, J., Munson, M., and Yoon, T.-Y. (2025) Exocyst stimulates multiple steps of exocytic SNARE complex assembly and vesicle fusion. Nat. Struct. Mol. Biol. 32, 150–160

18. Mei, K., Li, Y., Wang, S., Shao, G., Wang, J., Ding, Y., Luo, G., Yue, P., Liu, J.-J., Wang, X., Dong, M.-Q., Wang, H.-W., and Guo, W. (2018) Cryo-EM structure of the exocyst complex. Nat. Struct. Mol. Biol. 25, 139–146

19. Duan, Y., Guo, Q., Zhang, T., Meng, Y., Sun, D., Luo, G., and Liu, Y. (2019) Cyclin-dependent kinase– mediated phosphorylation of the exocyst subunit Exo84 in late G1 phase suppresses exocytic secretion and cell growth in yeast. J. Biol. Chem. 294, 11323–11332

20. Ren, J., and Guo, W. (2012) ERK1/2 Regulate Exocytosis through Direct Phosphorylation of the Exocyst Component Exo70. Dev. Cell. 22, 967–978

21. Chen, X.-W., Leto, D., Xiao, J., Goss, J., Wang, Q., Shavit, J. A., Xiong, T., Yu, G., Ginsburg, D., Toomre, D., Xu, Z., and Saltiel, A. R. (2011) Exocyst function regulated by effector phosphorylation. Nat. Cell Biol. 13, 580–588

22. Jin, R., Junutula, J. R., Matern, H. T., Ervin, K. E., Scheller, R. H., and Brunger, A. T. (2005) Exo84 and Sec5 are competitive regulatory Sec6/8 effectors to the RalA GTPase. EMBO J. 24, 2064–2074

23. Roumanie, O., Wu, H., Molk, J. N., Rossi, G., Bloom, K., and Brennwald, P. (2005) Rho GTPase regulation of exocytosis in yeast is independent of GTP hydrolysis and polarization of the exocyst complex. J. Cell Biol. 170, 583–594

24. Shelke, G. V., Williamson, C. D., Keren-Kaplan, T., and Bonifacino, J. S. (2026) TBC1D9 and TBC1D9B are ARL8 effectors that modulate exosome secretion through a RAB11A–exocyst pathway. Proc. Natl. Acad. Sci. 123, e2527864123

25. Guo, W., Roth, D., Gatti, E., De Camilli, P., and Novick, P. (1997) Identification and characterization of homologues of the Exocyst component Sec10p. FEBS Lett. 404, 135–139

26. Ting, A. E., Hazuka, C. D., Hsu, S. C., Kirk, M. D., Bean, A. J., and Scheller, R. H. (1995) rSec6 and rSec8, mammalian homologs of yeast proteins essential for secretion. Proc. Natl. Acad. Sci. 92, 9613– 9617

27. Fujimoto, B. A., Young, M., Carter, L., Pang, A. P. S., Corley, M. J., Fogelgren, B., and Polgar, N. (2019) The exocyst complex regulates insulin-stimulated glucose uptake of skeletal muscle cells. Am. J. Physiol.-Endocrinol. Metab. 317, E957–E972

28. Zuo, X., Kwon, S.-H., Janech, M. G., Dang, Y., Lauzon, S. D., Fogelgren, B., Polgar, N., and Lipschutz, J. H. (2019) Primary cilia and the exocyst are linked to urinary extracellular vesicle production and content. J. Biol. Chem. 294, 19099–19110

29. Zuo, X., Guo, W., and Lipschutz, J. H. (2009) The Exocyst Protein Sec10 Is Necessary for Primary Ciliogenesis and Cystogenesis In Vitro. Mol. Biol. Cell. 20, 2522–2529

30. Saint-Pol, J., and Culot, M. (2025) Minimum information for studies of extracellular vesicles (MISEV) as toolbox for rigorous, reproducible and homogeneous studies on extracellular vesicles. Toxicol. In Vitro. 106, 106049

31. Zhang, C., Brown, M. Q., Van De Ven, W., Zhang, Z.-M., Wu, B., Young, M. C., Synek, L., Borchardt, D., Harrison, R., Pan, S., Luo, N., Huang, Y. M., Ghang, Y.-J., Ung, N., Li, R., Isley, J., Morikis, D., Song, J., Guo, W., Hooley, R. J., Chang, C. A., Yang, Z., Zarsky, V., Muday, G. K., Hicks, G. R., and Raikhel, N. V. (2016) Endosidin2 targets conserved exocyst complex subunit EXO70 to inhibit exocytosis. Proc. Natl. Acad. Sci. 10.1073/pnas.1521248112

32. An, S. J., Anneken, A., Xi, Z., Choi, C., Schlessinger, J., and Toomre, D. (2022) Regulation of EGF-stimulated activation of the PI-3K/AKT pathway by exocyst-mediated exocytosis. Proc. Natl. Acad. Sci. 119, e2208947119

33. Newman, L. A., Useckaite, Z., and Rowland, A. (2022) Addressing MISEV guidance using targeted LC-MS/MS: A method for the detection and quantification of extracellular vesicle-enriched and contaminant protein markers from blood. J. Extracell. Biol. 1, e56

34. Ostrowski, M., Carmo, N. B., Krumeich, S., Fanget, I., Raposo, G., Savina, A., Moita, C. F., Schauer, K., Hume, A. N., Freitas, R. P., Goud, B., Benaroch, P., Hacohen, N., Fukuda, M., Desnos, C., Seabra, M. C., Darchen, F., Amigorena, S., Moita, L. F., and Thery, C. (2010) Rab27a and Rab27b control different steps of the exosome secretion pathway. Nat. Cell Biol. 12, 19–30

35. Sun, Q., Weng, R.-X., Li, J.-H., Li, Y.-C., Xu, J.-T., Li, R., Lu, X., and Xu, G.-Y. (2023) Rab27a-mediated exosome secretion in anterior cingulate cortex contributes to colorectal visceral pain in adult mice with neonatal maternal deprivation. Am. J. Physiol.-Gastrointest. Liver Physiol. 325, G356–G367

36. Song, L., Tang, S., Han, X., Jiang, Z., Dong, L., Liu, C., Liang, X., Dong, J., Qiu, C., Wang, Y., and Du, Y. (2019) KIBRA controls exosome secretion via inhibiting the proteasomal degradation of Rab27a. Nat. Commun. 10, 1639

37. Larijani, B., Hume, A. N., Tarafder, A. K., and Seabra, M. C. (2003) Multiple Factors Contribute to Inefficient Prenylation of Rab27a in Rab Prenylation Diseases. J. Biol. Chem. 278, 46798–46804

38. Arora, D. K., Syed, I., Machhadieh, B., McKenna, C. E., and Kowluru, A. (2012) Rab-geranylgeranyl transferase regulates glucose-stimulated insulin secretion from pancreatic β cells. Islets. 4, 354–358

39. Zhao, K., Matsunaga, K., Mizuno, K., Wang, H., Okunishi, K., and Izumi, T. (2023) Functional hierarchy among different Rab27 effectors involved in secretory granule exocytosis. eLife. 12, e82821

40. Chen, W., Kong, L., Connelly, S., Dendle, J. M., Liu, Y., Wilson, I. A., Powers, E. T., and Kelly, J. W. (2016) Stabilizing the C_H_ 2 Domain of an Antibody by Engineering in an Enhanced Aromatic Sequon. ACS Chem. Biol. 11, 1852–1861

41. Abramson, J., Adler, J., Dunger, J., Evans, R., Green, T., Pritzel, A., Ronneberger, O., Willmore, L., Ballard, A. J., Bambrick, J., Bodenstein, S. W., Evans, D. A., Hung, C.-C., O’Neill, M., Reiman, D., Tunyasuvunakool, K., Wu, Z., Žemgulytė, A., Arvaniti, E., Beattie, C., Bertolli, O., Bridgland, A., Cherepanov, A., Congreve, M., Cowen-Rivers, A. I., Cowie, A., Figurnov, M., Fuchs, F. B., Gladman, H., Jain, R., Khan, Y. A., Low, C. M. R., Perlin, K., Potapenko, A., Savy, P., Singh, S., Stecula, A., Thillaisundaram, A., Tong, C., Yakneen, S., Zhong, E. D., Zielinski, M., Žídek, A., Bapst, V., Kohli, P., Jaderberg, M., Hassabis, D., and Jumper, J. M. (2024) Accurate structure prediction of biomolecular interactions with AlphaFold 3. Nature. 630, 493–500

42. Sun, Y., Bilan, P. J., Liu, Z., and Klip, A. (2010) Rab8A and Rab13 are activated by insulin and regulate GLUT4 translocation in muscle cells. Proc. Natl. Acad. Sci. 107, 19909–19914

43. Essid, M., Gopaldass, N., Yoshida, K., Merrifield, C., and Soldati, T. (2012) Rab8a regulates the exocyst-mediated kiss-and-run discharge of the *Dictyostelium* contractile vacuole. Mol. Biol. Cell. 23, 1267–1282

44. Wu, S., Mehta, S. Q., Pichaud, F., Bellen, H. J., and Quiocho, F. A. (2005) Sec15 interacts with Rab11 via a novel domain and affects Rab11 localization in vivo. Nat. Struct. Mol. Biol. 12, 879–885

45. Li, X., Burnight, E. R., Cooney, A. L., Malani, N., Brady, T., Sander, J. D., Staber, J., Wheelan, S. J., Joung, J. K., McCray, P. B., Bushman, F. D., Sinn, P. L., and Craig, N. L. (2013) *piggyBac* transposase tools for genome engineering. Proc. Natl. Acad. Sci. 10.1073/pnas.1305987110

46. Johnson, E. T., Owens, J. B., and Moisyadi, S. (2016) Vast potential for using the *piggyBac* transposon to engineer transgenic plants at specific genomic locations. Bioengineered. 7, 3–6

47. Shin, S., Lee, S.-Y., Kang, M.-G., Jang, D.-G., Kim, J., Rhee, H.-W., and Kim, J.-S. (2024) Super-resolution proximity labeling with enhanced direct identification of biotinylation sites. *Commun*. Biol. 7, 554

48. Kubitz, L., Bitsch, S., Zhao, X., Schmitt, K., Deweid, L., Roehrig, A., Barazzone, E. C., Valerius, O., Kolmar, H., and Béthune, J. (2022) Engineering of ultraID, a compact and hyperactive enzyme for proximity-dependent biotinylation in living cells. *Commun*. Biol. 5, 657

49. Tarafder, A. K., Wasmeier, C., Figueiredo, A. C., Booth, A. E. G., Orihara, A., Ramalho, J. S., Hume, A. N., and Seabra, M. C. (2011) Rab27a Targeting to Melanosomes Requires Nucleotide Exchange but Not Effector Binding. Traffic. 12, 1056–1066

50. Fukuda, M. (2003) Slp4-a/Granuphilin-a Inhibits Dense-core Vesicle Exocytosis through Interaction with the GDP-bound Form of Rab27A in PC12 Cells. J. Biol. Chem. 278, 15390–15396

51. Bierings, R., Hellen, N., Kiskin, N., Knipe, L., Fonseca, A.-V., Patel, B., Meli, A., Rose, M., Hannah, M. J., and Carter, T. (2012) The interplay between the Rab27A effectors Slp4-a and MyRIP controls hormone-evoked Weibel-Palade body exocytosis. Blood. 120, 2757–2767

52. Liu, J., Zhao, Y., Sun, Y., He, B., Yang, C., Svitkina, T., Goldman, Y. E., and Guo, W. (2012) Exo70 Stimulates the Arp2/3 Complex for Lamellipodia Formation and Directional Cell Migration. Curr. Biol. 22, 1510–1515

53. Kee, A. J., Yang, L., Lucas, C. A., Greenberg, M. J., Martel, N., Leong, G. M., Hughes, W. E., Cooney, G. J., James, D. E., Ostap, E. M., Han, W., Gunning, P. W., and Hardeman, E. C. (2015) An Actin Filament Population Defined by the Tropomyosin Tpm3.1 Regulates Glucose Uptake. Traffic. 16, 691– 711

54. Desnos, C., Schonn, J.-S., Huet, S., Tran, V. S., El-Amraoui, A., Raposo, G., Fanget, I., Chapuis, C., Ménasché, G., De Saint Basile, G., Petit, C., Cribier, S., Henry, J.-P., and Darchen, F. (2003) Rab27A and its effector MyRIP link secretory granules to F-actin and control their motion towards release sites. J. Cell Biol. 163, 559–570

55. Wu, X., Wang, F., Rao, K., Sellers, J. R., and Hammer, J. A. (2002) Rab27a Is an Essential Component of Melanosome Receptor for Myosin Va. Mol. Biol. Cell. 13, 1735–1749

56. Sung, B. H., and Weaver, A. M. (2023) Visualization of Exosome Release and Uptake During Cell Migration Using the Live Imaging Reporter pHluorin_M153R-CD63. in Cell Migration in Three Dimensions (Margadant, C. ed), pp. 83–96, Methods in Molecular Biology, Springer US, New York, NY, 2608, 83–96

57. Sung, B. H., Von Lersner, A., Guerrero, J., Krystofiak, E. S., Inman, D., Pelletier, R., Zijlstra, A., Ponik, S. M., and Weaver, A. M. (2020) A live cell reporter of exosome secretion and uptake reveals pathfinding behavior of migrating cells. Nat. Commun. 11, 2092

58. Verweij, F. J., Revenu, C., Arras, G., Dingli, F., Loew, D., Pegtel, D. M., Follain, G., Allio, G., Goetz, J. G., Zimmermann, P., Herbomel, P., Del Bene, F., Raposo, G., and Van Niel, G. (2019) Live Tracking of Inter-organ Communication by Endogenous Exosomes In Vivo. Dev. Cell. 48, 573–589.e4

59. Zhao, F., Li, P., Chen, S. R. W., Louis, C. F., and Fruen, B. R. (2001) Dantrolene Inhibition of Ryanodine Receptor Ca2+Release Channels. J. Biol. Chem. 276, 13810–13816

60. Datta, A., Kim, H., Lal, M., McGee, L., Johnson, A., Moustafa, A. A., Jones, J. C., Mondal, D., Ferrer, M., and Abdel-Mageed, A. B. (2017) Manumycin A suppresses exosome biogenesis and secretion via targeted inhibition of Ras/Raf/ERK1/2 signaling and hnRNP H1 in castration-resistant prostate cancer cells. Cancer Lett. 408, 73–81

61. Bringas, C. F., Ahangar, M. S., Cuenco, J., Liu, H., Addinsall, A. B., Lindahl, M., Ovens, A. J., Febbraio, M. A., Foretz, M., Göransson, O., Scott, J. W., Zeqiraj, E., and Sakamoto, K. (2025) Mechanism and cellular actions of the potent AMPK inhibitor BAY-3827. Sci. Adv. 11, eadx2434

62. Calleja, V., Laguerre, M., Parker, P. J., and Larijani, B. (2009) Role of a Novel PH-Kinase Domain Interface in PKB/Akt Regulation: Structural Mechanism for Allosteric Inhibition. PLoS Biol. 7, e1000017

63. Chaikuad, A., M C Tacconi, E., Zimmer, J., Liang, Y., Gray, N. S., Tarsounas, M., and Knapp, S. (2014) A unique inhibitor binding site in ERK1/2 is associated with slow binding kinetics. Nat. Chem. Biol. 10, 853–860

64. Wong, M. H., Samal, A. B., Lee, M., Vlach, J., Novikov, N., Niedziela-Majka, A., Feng, J. Y., Koltun, D. O., Brendza, K. M., Kwon, H. J., Schultz, B. E., Sakowicz, R., Saad, J. S., and Papalia, G. A. (2019) The KN-93 Molecule Inhibits Calcium/Calmodulin-Dependent Protein Kinase II (CaMKII) Activity by Binding to Ca2+/CaM. J. Mol. Biol. 431, 1440–1459

65. Gschwendt, M., Dieterich, S., Rennecke, J., Kittstein, W., Mueller, H.-J., and Johannes, F.-J. (1996) Inhibition of protein kinase C μ by various inhibitors. Inhibition from protein kinase c isoenzymes. FEBS Lett. 392, 77–80

66. Mu, J., Brozinick, J. T., Valladares, O., Bucan, M., and Birnbaum, M. J. (2001) A Role for AMP-Activated Protein Kinase in Contraction- and Hypoxia-Regulated Glucose Transport in Skeletal Muscle. Mol. Cell. 7, 1085–1094

67. Boyer, J. G., Prasad, V., Song, T., Lee, D., Fu, X., Grimes, K. M., Sargent, M. A., Sadayappan, S., and Molkentin, J. D. (2019) ERK1/2 signaling induces skeletal muscle slow fiber-type switching and reduces muscular dystrophy disease severity. JCI Insight. 4, e127356

68. Parkington, J. D., LeBrasseur, N. K., Siebert, A. P., and Fielding, R. A. (2004) Contraction-mediated mTOR, p70^S^6^k^, and ERK1/2 phosphorylation in aged skeletal muscle. J. Appl. Physiol. 97, 243–248

69. Kramer, H. F., Witczak, C. A., Taylor, E. B., Fujii, N., Hirshman, M. F., and Goodyear, L. J. (2006) AS160 Regulates Insulin- and Contraction-stimulated Glucose Uptake in Mouse Skeletal Muscle. J. Biol. Chem. 281, 31478–31485

70. Nakanishi, A., Hatano, N., Fujiwara, Y., Sha’ri, A., Takabatake, S., Akano, H., Kanayama, N., Magari, M., Nozaki, N., and Tokumitsu, H. (2017) AMP-activated protein kinase–mediated feedback phosphorylation controls the Ca2+/calmodulin (CaM) dependence of Ca2+/CaM-dependent protein kinase kinase β. J. Biol. Chem. 292, 19804–19813

71. Schmitt, J. M., Wayman, G. A., Nozaki, N., and Soderling, T. R. (2004) Calcium Activation of ERK Mediated by Calmodulin Kinase I. J. Biol. Chem. 279, 24064–24072

72. Roskoski, R. (2012) ERK1/2 MAP kinases: Structure, function, and regulation. Pharmacol. Res. 66, 105– 143

73. Eguez, L., Lee, A., Chavez, J. A., Miinea, C. P., Kane, S., Lienhard, G. E., and McGraw, T. E. (2005) Full intracellular retention of GLUT4 requires AS160 Rab GTPase activating protein. Cell Metab. 2, 263–272

74. Kang, J. K., Kim, O.-H., Hur, J., Yu, S. H., Lamichhane, S., Lee, J. W., Ojha, U., Hong, J. H., Lee, C. S., Cha, J.-Y., Lee, Y. J., Im, S.-S., Park, Y. J., Choi, C. S., Lee, D. H., Lee, I.-K., and Oh, B.-C. (2017) Increased intracellular Ca^2+^ concentrations prevent membrane localization of PH domains through the formation of Ca^2+^-phosphoinositides. Proc. Natl. Acad. Sci. 114, 11926–11931

75. Guo, L., Urban, J. F., Zhu, J., and Paul, W. E. (2008) Elevating Calcium in Th2 Cells Activates Multiple Pathways to Induce IL-4 Transcription and mRNA Stabilization. J. Immunol. 181, 3984–3993

76. Park, C. H., Kim, Y. S., Kim, Y. H., Choi, M. Y., Yoo, J. M., Kang, S. S., Choi, W. S., and Cho, G. J. (2008) Calcineurin mediates AKT dephosphorylation in the ischemic rat retina. Brain Res. 1234, 148–157

77. Treebak, J. T., Glund, S., Deshmukh, A., Klein, D. K., Long, Y. C., Jensen, T. E., Jørgensen, S. B., Viollet, B., Andersson, L., Neumann, D., Wallimann, T., Richter, E. A., Chibalin, A. V., Zierath, J. R., and Wojtaszewski, J. F. P. (2006) AMPK-Mediated AS160 Phosphorylation in Skeletal Muscle Is Dependent on AMPK Catalytic and Regulatory Subunits. Diabetes. 55, 2051–2058

78. Tan, S.-X., Ng, Y., Burchfield, J. G., Ramm, G., Lambright, D. G., Stöckli, J., and James, D. E. (2012) The Rab GTPase-Activating Protein TBC1D4/AS160 Contains an Atypical Phosphotyrosine-Binding Domain That Interacts with Plasma Membrane Phospholipids To Facilitate GLUT4 Trafficking in Adipocytes. Mol. Cell. Biol. 32, 4946–4959

79. Williams, J. K., Ngo, J. M., Lehman, I. M., and Schekman, R. (2023) Annexin A6 mediates calcium-dependent exosome secretion during plasma membrane repair. eLife. 12, e86556

80. Pandey, M. P., Pezeshkian, W., and Khandelia, H. (2023) Annexin proteins involved in plasma membrane repair reshape a membrane edge: Direct evidence from molecular simulations. Biophys. J. 122, 323a

81. Boye, T. L., Maeda, K., Pezeshkian, W., Sønder, S. L., Haeger, S. C., Gerke, V., Simonsen, A. C., and Nylandsted, J. (2017) Annexin A4 and A6 induce membrane curvature and constriction during cell membrane repair. Nat. Commun. 8, 1623

82. Wu, H., Nguyen, H., Hashim, P. H., Fogelgren, B., Duncan, F. E., and Ward, W. S. (2024) Oocyte-specific EXOC5 expression is required for mouse oogenesis and folliculogenesis. Mol. Hum. Reprod. 30, gaae026

83. Lee, B., Baek, J.-I., Min, H., Bae, S.-H., Moon, K., Kim, M.-A., Kim, Y.-R., Fogelgren, B., Lipschutz, J. H., Lee, K.-Y., Bok, J., and Kim, U.-K. (2018) Exocyst Complex Member EXOC5 Is Required for Survival of Hair Cells and Spiral Ganglion Neurons and Maintenance of Hearing. Mol. Neurobiol. 55, 6518–6532

84. Thapa, N., Sun, Y., Schramp, M., Choi, S., Ling, K., and Anderson, R. A. (2012) Phosphoinositide Signaling Regulates the Exocyst Complex and Polarized Integrin Trafficking in Directionally Migrating Cells. Dev. Cell. 22, 116–130

85. Rameh, L. E., Tolias, K. F., Duckworth, B. C., and Cantley, L. C. (1997) A new pathway for synthesis of phosphatidylinositol-4,5-bisphosphate. Nature. 390, 192–196

86. Park, S. J., Itoh, T., and Takenawa, T. (2001) Phosphatidylinositol 4-Phosphate 5-Kinase Type I Is Regulated through Phosphorylation Response by Extracellular Stimuli. J. Biol. Chem. 276, 4781–4787

87. Liu, A., Sui, D., Wu, D., and Hu, J. (2016) The activation loop of PIP5K functions as a membrane sensor essential for lipid substrate processing. Sci. Adv. 2, e1600925

88. Liu, J., Zuo, X., Yue, P., and Guo, W. (2007) Phosphatidylinositol 4,5-Bisphosphate Mediates the Targeting of the Exocyst to the Plasma Membrane for Exocytosis in Mammalian Cells. Mol. Biol. Cell. 18, 4483–4492

89. Wang, J., Ren, J., Wu, B., Feng, S., Cai, G., Tuluc, F., Peränen, J., and Guo, W. (2015) Activation of Rab8 guanine nucleotide exchange factor Rabin8 by ERK1/2 in response to EGF signaling. Proc. Natl. Acad. Sci. 112, 148–153

90. Wang, B., Liang, Z., Tan, T., Zhang, M., Jiang, Y., Shang, Y., Gao, X., Song, S., Wang, R., Chen, H., Liu, J., Li, J., Ren, Y., and Liu, P. (2023) CRB3 navigates Rab11 trafficking vesicles to promote γTuRC assembly during ciliogenesis. eLife. 12, RP86689

91. Prekeris, R., Klumperman, J., and Scheller, R. H. (2000) A Rab11/Rip11 Protein Complex Regulates Apical Membrane Trafficking via Recycling Endosomes. Mol. Cell. 6, 1437–1448

92. Biesemann, A., Gorontzi, A., Barr, F., and Gerke, V. (2017) Rab35 protein regulates evoked exocytosis of endothelial Weibel–Palade bodies. J. Biol. Chem. 292, 11631–11640

93. Guo, D., Tikoo, S., and Haass, N. K. (2019) RAB27A-mediated melanoma exosomes: promoters of invasion and metastasis. Transl. Cancer Res. 8, 732–735

94. Li, W., Hu, Y., Jiang, T., Han, Y., Han, G., Chen, J., and Li, X. (2014) Rab27A regulates exosome secretion from lung adenocarcinoma cells A549: involvement of EPI 64. APMIS. 122, 1080–1087

95. Rinaldi, F. C., Packer, M., and Collins, R. (2015) New insights into the molecular mechanism of the Rab GTPase Sec4p activation. BMC Struct. Biol. 15, 14

96. Ramalho, J. S., Anders, R., Jaissle, G. B., Seeliger, M. W., Huxley, C., and Seabra, M. C. (2002) Rapid degradation of dominant-negative Rab27 proteins in vivo precludes their use in transgenic mouse models. BMC Cell Biol. 3, 26

97. Goler-Baron, V., Sladkevich, I., and Assaraf, Y. G. (2012) Inhibition of the PI3K-Akt signaling pathway disrupts ABCG2-rich extracellular vesicles and overcomes multidrug resistance in breast cancer cells. Biochem. Pharmacol. 83, 1340–1348

98. Wong, M. H., Samal, A. B., Lee, M., Vlach, J., Novikov, N., Niedziela-Majka, A., Feng, J. Y., Koltun, D. O., Brendza, K. M., Kwon, H. J., Schultz, B. E., Sakowicz, R., Saad, J. S., and Papalia, G. A. (2019) The KN-93 Molecule Inhibits Calcium/Calmodulin-Dependent Protein Kinase II (CaMKII) Activity by Binding to Ca2+/CaM. J. Mol. Biol. 431, 1440–1459

99. Miranda, L., Carpentier, S., Platek, A., Hussain, N., Gueuning, M.-A., Vertommen, D., Ozkan, Y., Sid, B., Hue, L., Courtoy, P. J., Rider, M. H., and Horman, S. (2010) AMP-activated protein kinase induces actin cytoskeleton reorganization in epithelial cells. Biochem. Biophys. Res. Commun. 396, 656–661

100. Shrestha, M. M., Lim, C.-Y., Bi, X., Robinson, R. C., and Han, W. (2021) Tmod3 Phosphorylation Mediates AMPK-Dependent GLUT4 Plasma Membrane Insertion in Myoblasts. Front. Endocrinol. 12, 653557

101. Mitani, F., Lin, J., Sakamoto, T., Uehara, R., Hikita, T., Yoshida, T., Setiawan, A., Arai, M., and Oneyama, C. (2022) Asteltoxin inhibits extracellular vesicle production through AMPK/mTOR-mediated activation of lysosome function. Sci. Rep. 12, 6674

102. Zhang, T., Sun, D., Luo, G., and Liu, Y. (2019) Spatial and Translational Regulation of Exocyst Subunits by Cell Cycle in Budding Yeast. Med. Sci. Monit. 25, 4059–4067

103. Honorato, R. V., Trellet, M. E., Jiménez-García, B., Schaarschmidt, J. J., Giulini, M., Reys, V., Koukos, P. I., Rodrigues, J. P. G. L. M., Karaca, E., Van Zundert, G. C. P., Roel-Touris, J., Van Noort, C. W., Jandová, Z., Melquiond, A. S. J., and Bonvin, A. M. J. J. (2024) The HADDOCK2.4 web server for integrative modeling of biomolecular complexes. Nat. Protoc. 19, 3219–3241

104. Honorato, R. V., Koukos, P. I., Jiménez-García, B., Tsaregorodtsev, A., Verlato, M., Giachetti, A., Rosato, A., and Bonvin, A. M. J. J. (2021) Structural Biology in the Clouds: The WeNMR-EOSC Ecosystem. Front. Mol. Biosci. 8, 729513

105. Jiménez-García, B., Elez, K., Koukos, P. I., Bonvin, A. M., and Vangone, A. (2019) PRODIGY-crystal: a web-tool for classification of biological interfaces in protein complexes. Bioinformatics. 35, 4821–4823

106. Wiśniewski, J. R., Zougman, A., Nagaraj, N., and Mann, M. (2009) Universal sample preparation method for proteome analysis. Nat. Methods. 6, 359–362

107. Graw, S., Tang, J., Zafar, M. K., Byrd, A. K., Bolden, C., Peterson, E. C., and Byrum, S. D. (2020) proteiNorm – A User-Friendly Tool for Normalization and Analysis of TMT and Label-Free Protein Quantification. ACS Omega. 5, 25625–25633

108. Thurman, T. J., Washam, C. L., Alkam, D., Bird, J. T., Gies, A., Dhusia, K., Ii, M. S. R., and Byrum, S. D. (2023) proteoDA: a package for quantitative proteomics. J. Open Source Softw. 8, 5184

109. R Core Team (2026) R: A Language and Environment for Statistical Computing

110. Wickham, H., Averick, M., Bryan, J., Chang, W., McGowan, L., François, R., Grolemund, G., Hayes, A., Henry, L., Hester, J., Kuhn, M., Pedersen, T., Miller, E., Bache, S., Müller, K., Ooms, J., Robinson, D., Seidel, D., Spinu, V., Takahashi, K., Vaughan, D., Wilke, C., Woo, K., and Yutani, H. (2019) Welcome to the Tidyverse. J. Open Source Softw. 4, 1686

111. Wickham, H. (2009) ggplot2: Elegant Graphics for Data Analysis, Springer New York, New York, NY, 10.1007/978-0-387-98141-3

