## Supplementary material for "Calcium signaling recruits Rab27a and the exocyst complex to promote small extracellular vesicle release in skeletal muscle cells": Supp S1-10

**SUPPLEMENTAL FIGURES**

**Supp. S1**

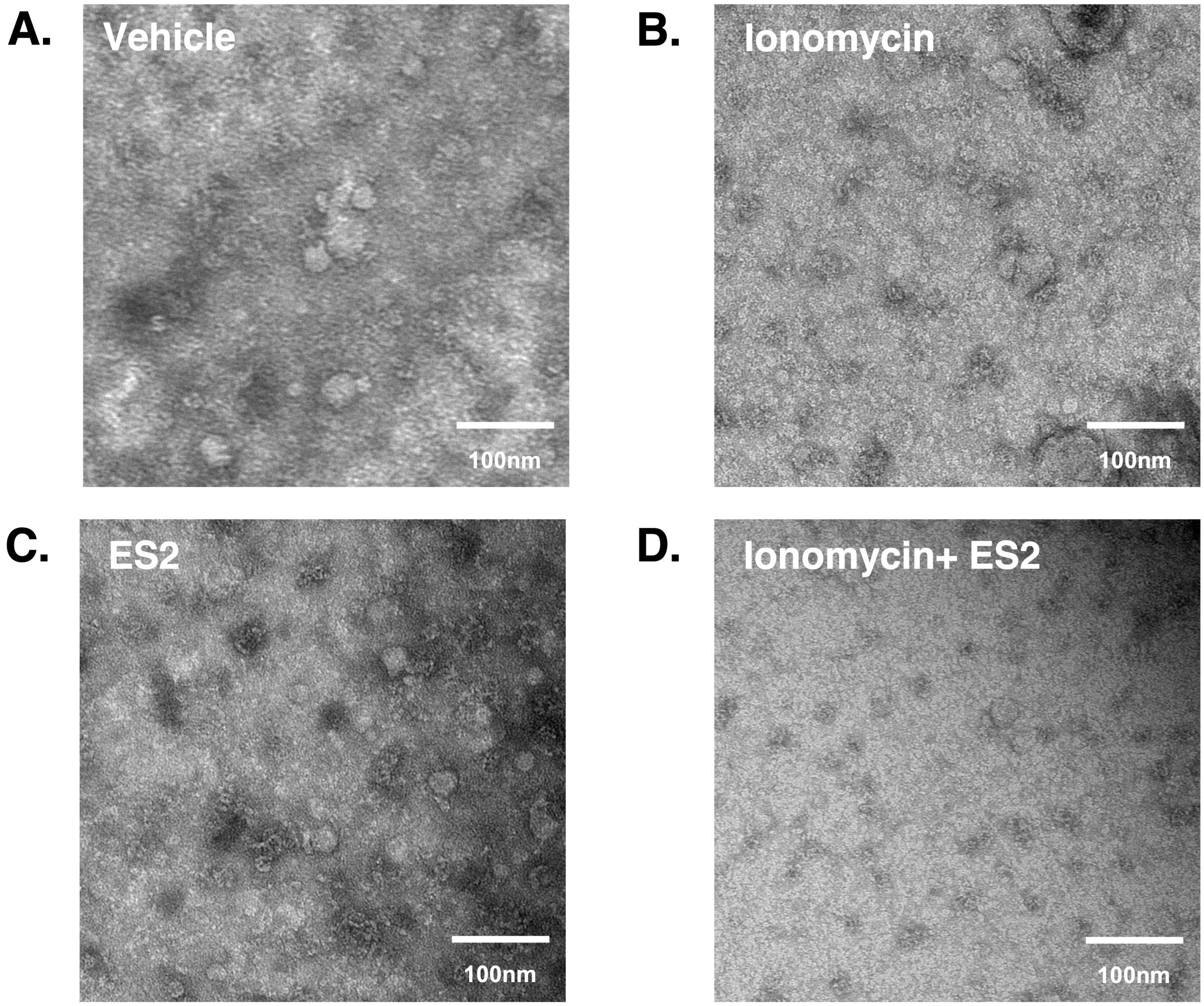

**Supp. S2**

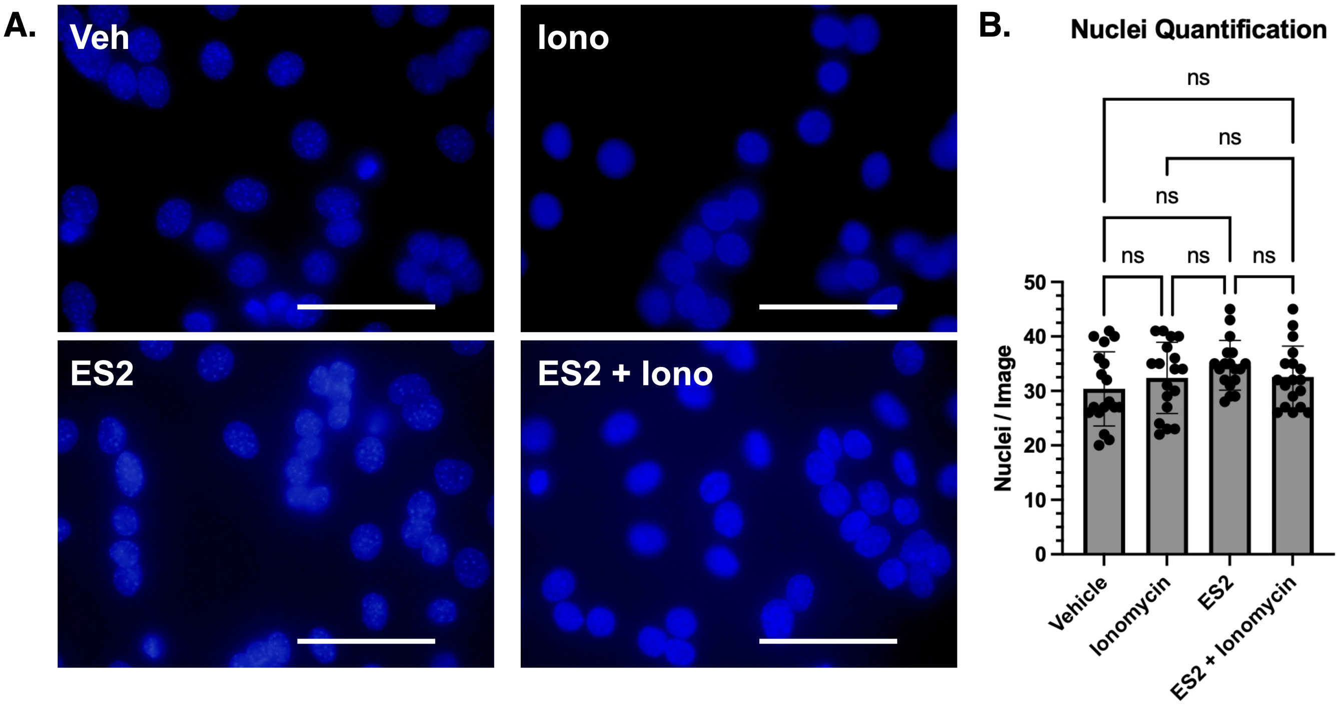
A.
Veh
Iono
ES2
ES2 + Iono
B.
Nuclei Quantification
ns
ns
ns
ns
ns
ns
50
40
30
20
10
0
Nuclei / Image
Vehicle
Ionomycin
ES2
ES2 + Ionomycin

### Supp. S3

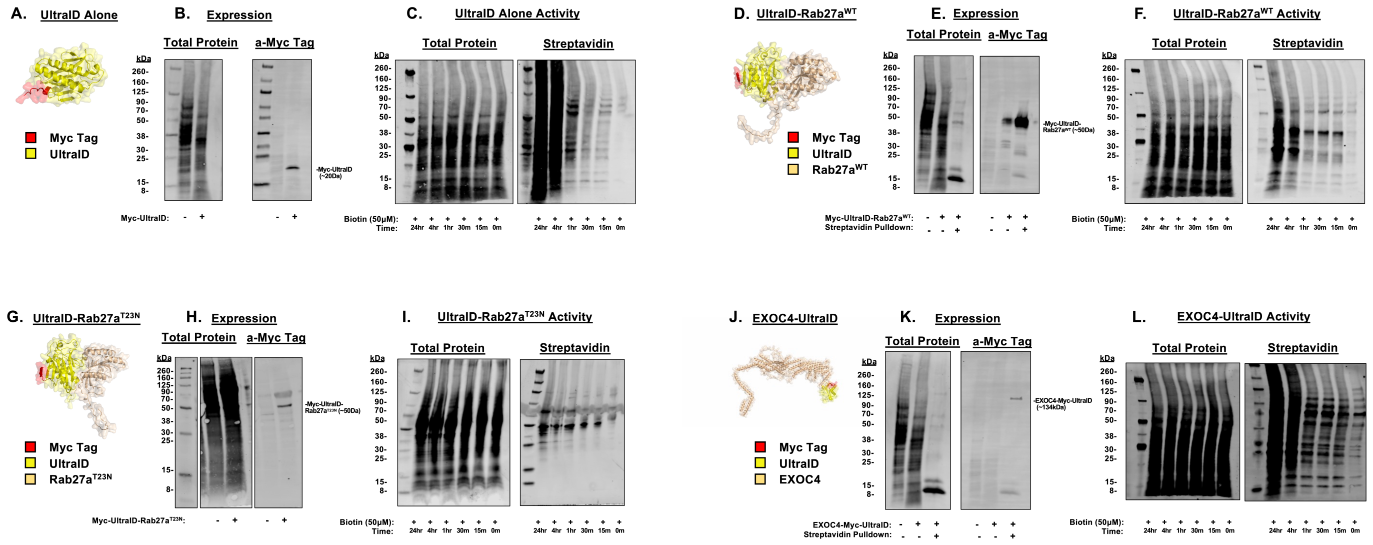

A. UltraID Alone
Myc Tag
UltraID
B. Expression
Total Protein
a-Myc Tag
kDa
Myc-UltraID:
C. UltraID Alone Activity
Total Protein
Streptavidin
Biotin (50μM):
Time:
24hr 4hr 1hr 30m 15m 0m
D. UltraID-Rab27a^WT
Myc Tag
UltraID
Rab27a^WT
E. Expression
Total Protein
a-Myc Tag
Myc-UltraID-Rab27a^WT:
Streptavidin Pulldown:
F. UltraID-Rab27a^WT Activity
Total Protein
Streptavidin
G. UltraID-Rab27a^T23N
Myc Tag
UltraID
Rab27a^T23N
H. Expression
Total Protein
a-Myc Tag
Myc-UltraID-Rab27a^T23N:
I. UltraID-Rab27a^T23N Activity
Total Protein
Streptavidin
J. EXOC4-UltraID
Myc Tag
UltraID
EXOC4
K. Expression
Total Protein
a-Myc Tag
EXOC4-Myc-UltraID:
Streptavidin Pulldown:
L. EXOC4-UltraID Activity
Total Protein
Streptavidin

**Supp. S4**

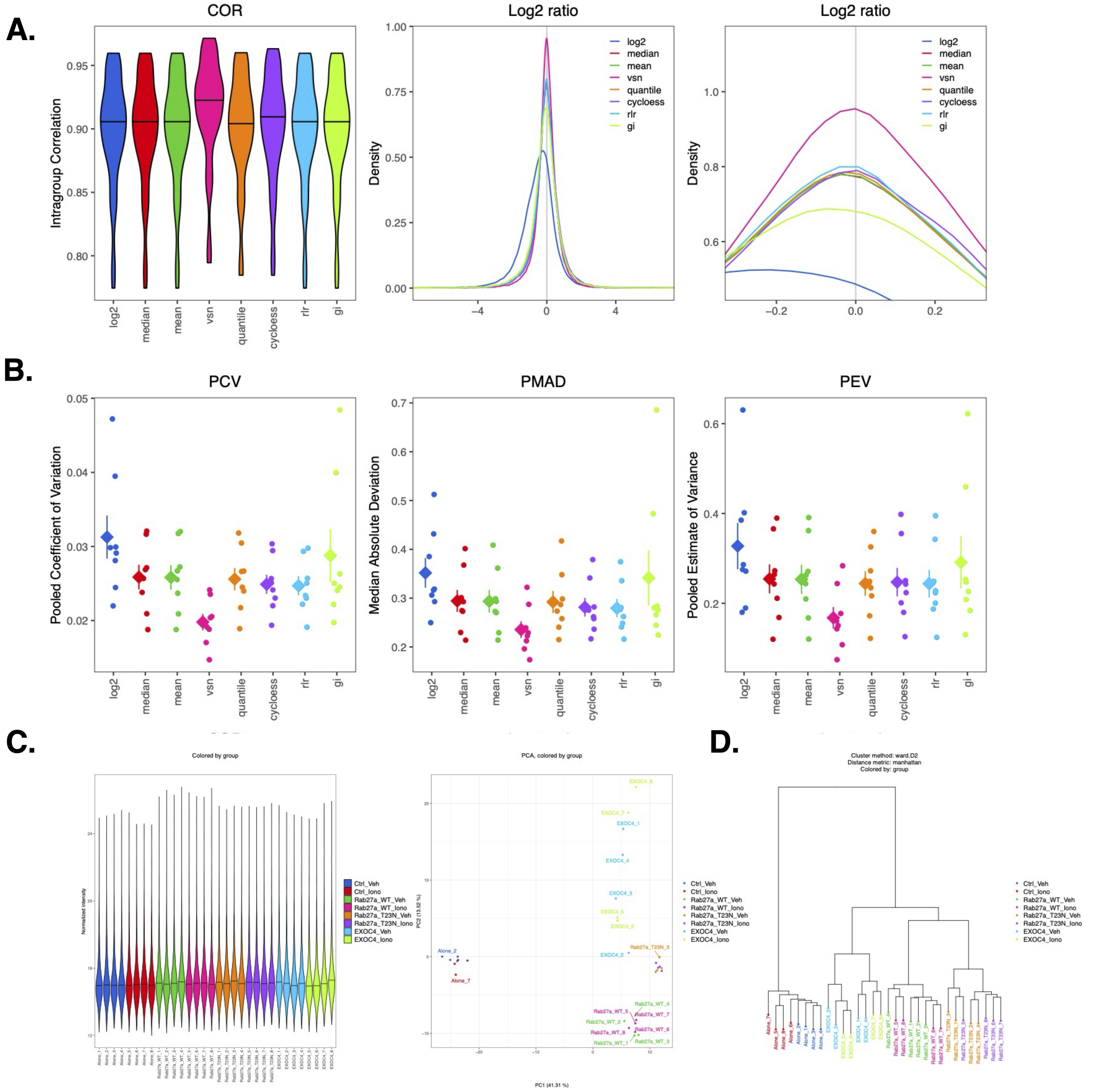

A.
COR
Intragroup Correlation
log2
median
mean
vsn
quantile
cycloess
rlr
gi
Log2 ratio
Density
Log2 ratio
Density
B.
PCV
Pooled Coefficient of Variation
PMAD
Median Absolute Deviation
PEV
Pooled Estimate of Variance
C.
Colored by group
Normalized intensity
Ctrl_Veh
Ctrl_Iono
Rab27a_WT_Veh
Rab27a_WT_Iono
Rab27a_T23N_Veh
Rab27a_T23N_Iono
EXOC4_Veh
EXOC4_Iono
PCA, colored by group
PC1 (41.31 %)
D.
Cluster method: ward.D2
Distance metric: manhattan
Colored by: group

Supp. S5

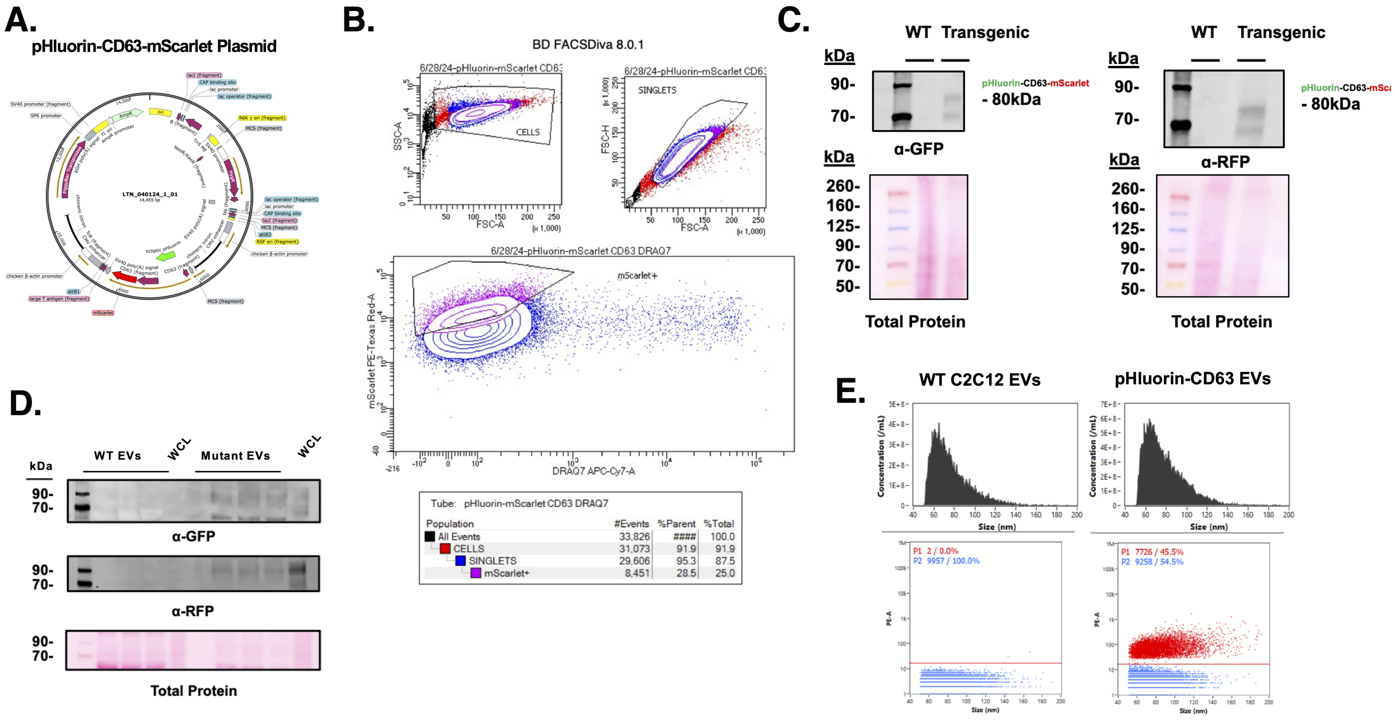

A.
pHluorin-CD63-mScarlet Plasmid
B.
BD FACSDiva 8.0.1
CELLS
SINGLETS
mScarlet+
Tube: pHluorin-mScarlet CD63 DRAQ7
Population #Events %Parent %Total
All Events 33,826 #### 100.0
CELLS 31,073 91.9 91.9
SINGLETS 29,606 95.3 87.5
mScarlet+ 8,451 28.5 25.0
C.
WT Transgenic
kDa
90-
70-
- 80kDa
α-GFP
α-RFP
260-
160-
125-
50-
Total Protein
D.
WT EVs
WCL
Mutant EVs
α-GFP
α-RFP
Total Protein
E.
WT C2C12 EVs
pHluorin-CD63 EVs
Concentration (/mL)
Size (nm)
P1 2 / 0.0%
P2 9957 / 100.0%
P1 7726 / 45.5%
P2 9258 / 54.5%

**Supp. S6**

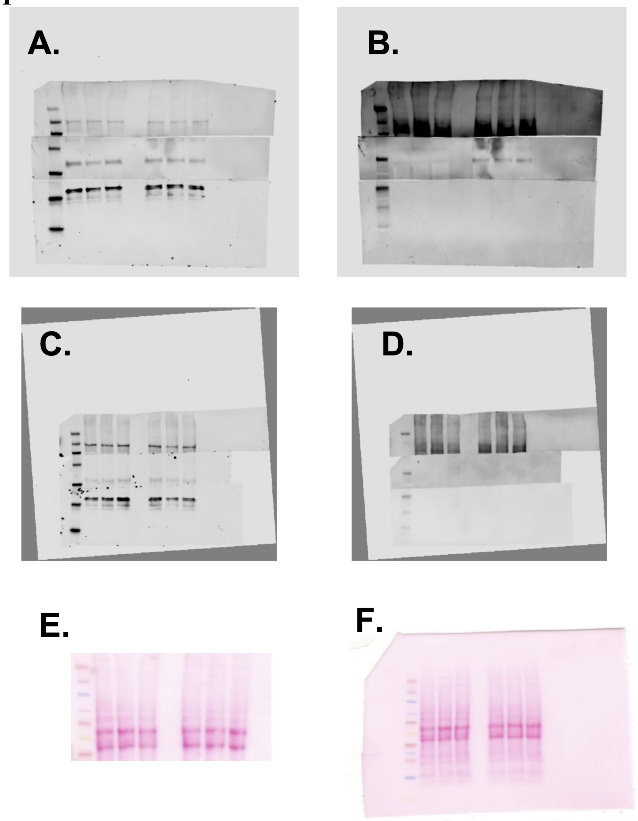

**Supp. S7**

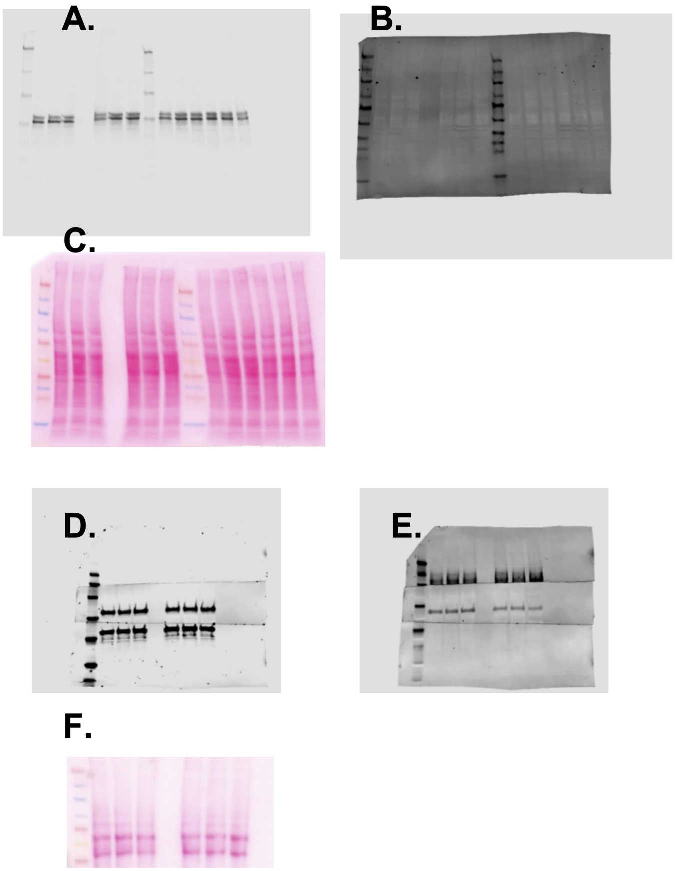

**Supp. S8**

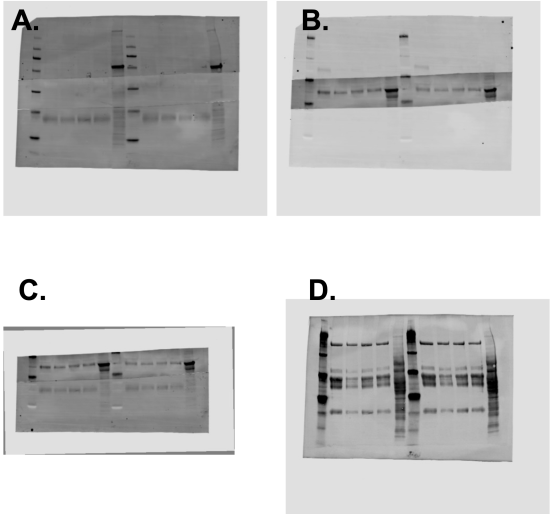

**Supp. S9**

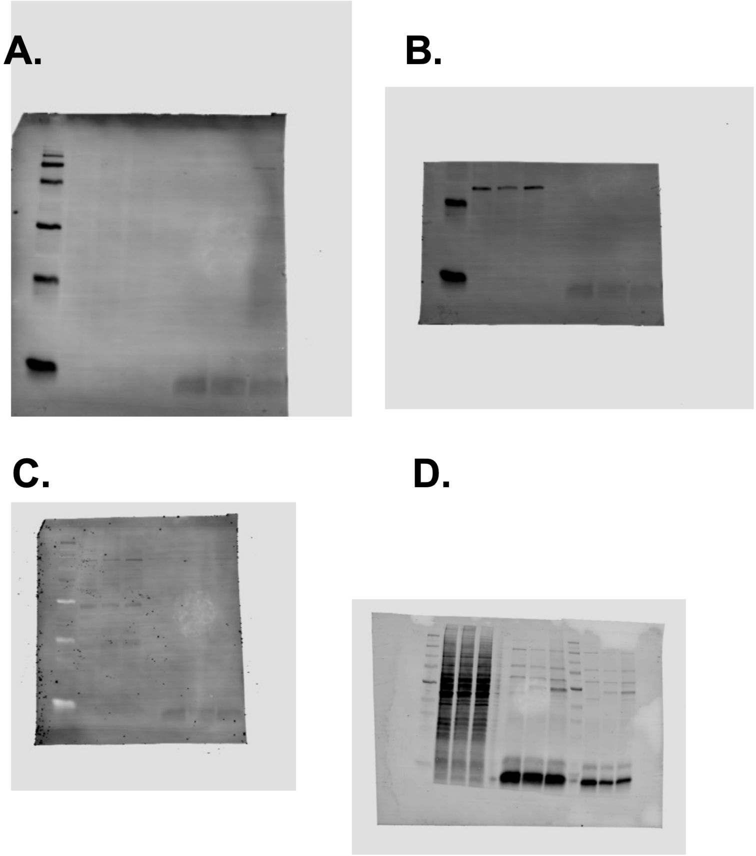

**Supp. S10**

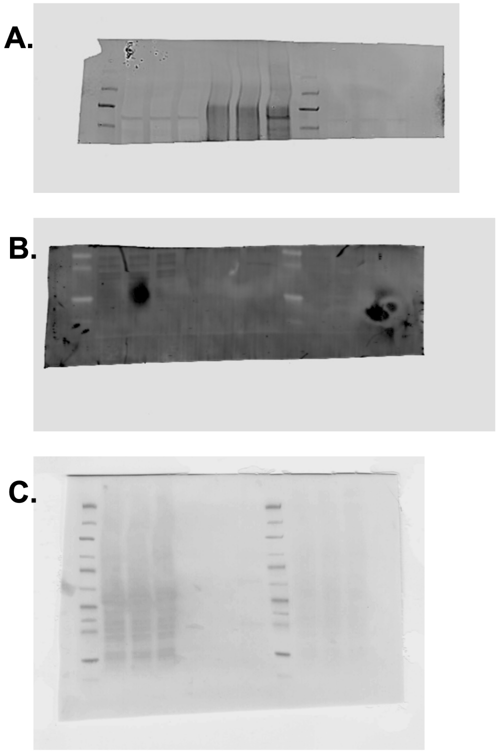

### SUPPLEMENTAL FIGURE LEGENDS

**Supplemental Figure S1. Representative Transmission Electron Microscopy Images**. TEM images of EVs isolated during treatment of C2C12 myotubes with A. Vehicle, B. Ionomycin, C. ES2, and D. Ionomycin + ES2 (corresponding to **Fig. 2**).

**Supplemental Figure S2. Nuclei Quantification of EV-Producing Myotubes**. A. Epifluorescent representative images of C2C12 myotubes used during treatment and harvest of EVs (corresponding to **Fig. 2**). B. Quantification of nuclei in field of view (scale bar=50 µm) in myotubes treated with Vehicle, Ionomycin, ES2 and ES2 + Ionomycin (One-Way ANOVA; n=18).

**Supplemental Figure S3. Validating UltraID expressing SkM cell lines**. A-C. Predicted structure, expression (Myc Tag) and activity of Myc-UltraID in C2C12 myotubes (corresponding to **Fig. 8**). D-F. Predicted structure, expression (Myc Tag) and activity of Myc-UltraID-Rab27a$^{WT}$ in C2C12 myotubes. G-I. Predicted structure, expression (Myc Tag) and activity of Myc-UltraID-Rab27a$^{T23N}$ in C2C12 myotubes. J-L. Predicted structure, expression (Myc Tag) and activity of EXOC4-Myc-UltraID in C2C12 myotubes. Western blot membranes were probed with fluorescent Streptavidin$_{800\ nm}$ to resolve biotinylation levels over time (0-24hr) upon treatment with biotin (50 µm).

**Supplemental Figure S4. Quality control for LMCS proteomics**. A. Normalization methods including intragroup correlation coefficients (COR) and density distributions of log$_2$-transformed protein abundance ratios are shown for each normalization approach (corresponding to **Fig. 8**). B. Variance metrics across normalization methods, including pooled coefficient of variation (PCV), median absolute deviation (PMAD), and pooled estimate of variance (PEV). C. Global assessment of sample distributions and variance structure. Left: violin plots of normalized protein intensities across all samples, demonstrating comparable intensity distributions. Right: principal component analysis (PCA) of protein abundances, colored by experimental group, showing clustering of biological replicates and separation between conditions. D. Hierarchical clustering of samples based on protein abundance profiles using Ward's method and Manhattan distance.

**Supplemental Figure S5. Creating a stable expressing, monoclonal pHluorin-CD63 SkM line**. Methodology and materials behind creation of pHluorin-CD63-mScarlet monoclonal SkM cell line (corresponding to **Fig. 9**). A. Whole plasmid sequence result of pHluorin-CD63-mScarlet plasmid featuring piggyBac transposase gene and hygromycin antibiotic resistance gene. B. Schematic of flow cytometry gates for mScarlet positive, mScarlet negative, and viable (DRAQ7) C2C12 cells used to isolate single cells. C. Western blot of WT C2C12 vs pHluorin-CD63-mScarlet monoclonal C2C12 cell lysates probed with α-pan RFP and α-pan GFP antibodies. D. Western blotting of WT EVs and lysate vs pHluorin-CD63-mScarlet EVs and lysate. E. Nano flow cytometry of unlabeled EVs produced from WT C2C12 myotubes (left) vs EVs produced from pHluorin-CD63-mScarlet monoclonal C2C12 myotubes (right) taken in PE-A red fluorescent channel (540/80 nm).

**Supplemental Figure S6. Uncropped Western Blots I**. Uncropped western blots for kinases, substrates and phosphorylated kinases and substrates . A. Membrane probed for AS160 (top), AMPKa (middle), and GAPDH (bottom) (corresponding to **Fig. 1C**). B. Membrane probed for pAS160 (T642) (top) and pAMPK (T172) (middle) (corresponding to **Fig. 1C,D**). C. Membrane probed for AS160 (top), AMPKa (middle), and GAPDH (bottom) (corresponding to **Fig. 1C,D**). D. Membrane probed for pAS160 (T642) (corresponding to **Fig. 1D**). E. Total protein stain for AMPK and AS160 blots (A-B) (corresponding to **Fig. 1C,D**). F. Total protein stain for AMPK and AS160 blots (C-D) (corresponding to **Fig. 1C,D**).

**Supplemental Figure S7. Uncropped Western Blots II**. Uncropped western blots for kinases, substrates and phosphorylated kinases and substrates cont.'. A. Membrane probed for a-ERK1/2 (corresponding to **Fig. 1E**). B. Membrane probed for pERK1/2 (T202/T204) (corresponding to **Fig. 1E**). C. Total protein stain for ERK1/2 and pERK blots (corresponding to **Fig. 1E**). D. Membrane probed for AKT (top) and GAPDH (middle) (corresponding to **Fig. 1F**). E. Membrane probed for pAS160 (T642) (top) and pAKT (S473) (middle). F. Total protein stain for AKT and AS160 blots (D-E) (corresponding to **Fig. 1F**).

**Supplemental Figure S8. Uncropped Western Blots II**. Uncropped western blots for EV and cellular markers. A. Membrane probed for Calnexin (top) and CD81 (bottom). B. Membrane probed for TSG101 (middle). C. Membrane probed for TSG101 (middle) and CD9 (bottom). D. Total protein stain for EV and cellular lysate membrane (A-C) (corresponding to **Fig. 2B**).

**Supplemental Figure S9. Uncropped Western Blots III**. Uncropped western blots for Co-IP. A. Membrane probed for EXOC4 (SEC8). B. Rab27a. C. Total protein stain for Co-IP inputs and IP lanes (A-B) (corresponding to **Fig. 6A**).

**Supplemental Figure S10. Uncropped Western Blots IV**. Uncropped western blots for UltraID-Rab27aWT pulldown. A. Membrane probed for SYTL4. B. Membrane probed for GAPDH. C. Membrane probed for EXOC4 (SEC8). D. Total protein stain for UltraID-Rab27a Pulldown (A-C) (corresponding to **Fig. 8B**).
